# Dopaminergic modulation of behavioral and electrocortical markers of interpersonal performance monitoring in Parkinson’s Disease: insights from multivariate and univariate analyses

**DOI:** 10.1101/2024.08.14.607937

**Authors:** V. Era, U.G. Pesci, Q. Moreau, R. Pezzetta, S. Zabberoni, A. Peppe, A. Costa, S. Taglieri, M. Candidi, S.M. Aglioti

## Abstract

Effective interpersonal interaction necessitates constant monitoring and adaptation to others’ actions, a process known as interpersonal performance monitoring, which is influenced by the dopaminergic system and marked by specific electrocortical signatures. To explore the connection between deficits in interpersonal performance monitoring and altered neural markers, we assessed patients with Parkinson’s Disease (PD) performing coordination tasks with a virtual partner (VP) under two conditions: on dopaminergic medication (PD ON) and after withdrawal (PD OFF). In Interactive trials, which required adaptation to the VP’s actions, PD OFF performance was impaired compared to PD ON. EEG analysis revealed in PD OFF increased midfrontal Delta-Theta activity during Interactive trials. Higher Delta-Theta synchronization was associated with improved performance, suggesting compensatory mechanisms. Multivariate EEG analysis distinguished Interactive from Cued trials, especially in PD OFF. Our findings highlight dopamine’ s role in modulating electrocortical markers of interpersonal performance monitor, with significant implications for understanding and treating PD.

## INTRODUCTION

The gradual degeneration of dopaminergic neurons in the substantia nigra pars compacta, a hallmark of Parkinson’s disease (PD), affects the functioning of subcortical and cortical areas, especially frontal and cingulate regions (Ullsperger & Von Cramon, 2006; Wylie et al., 2010). This cascade of changes has been shown to lead to motor symptoms and impairments in higher-order cognitive functions (Blesa et al., 2022; Ponsi et al., 2021). A prominent feature characterizing cognitive and motor domains in PD is inflexible behaviour, marked by difficulties in adapting to external requirements when necessary (Galea et al., 2012). Behavioural flexibility is essential for effectively coordinating actions with a partner during everyday interactions and purposeful joint actions, demanding the capacity to predict and adaptively respond to the actions of others (Sebanz et al., 2006). Indeed, interpersonal interactions can go wrong in many ways, from communication breakdowns to conflicts and lack of reciprocity, underscoring the critical need for continuous monitoring of others’ behaviour, referred to as interpersonal performance monitoring (Era et al., 2019). Examining unexpected events during joint actions thus offers valuable insights into how the performance monitoring system allocates resources to oversee others’ actions. Behavioural studies reveal that processes resembling those engaged during self-generated errors also come into play when witnessing another person’s mistakes and when interacting with others in response to their errors (Sacheli et al., 2021). Furthermore, electrocortical signatures associated with error processing (i.e., increase in midfrontal theta power, the error-related negativity-ERN, the early and late error positivity, Pe) reflecting the neurocomputational processing of the monitoring system during individual motor tasks (Cohen, 2011; Falkenstein et al., 1991, 2000 Luu et al., 2004; Fusco et al., 2019; 2022) are also elicited when interacting with others in response to their errors or in conditions necessitating continuous monitoring and adaptation to their actions (Era et al., 2019; Moreau et al., 2020, 2022). Moreover, studies using non-invasive brain stimulation have shown that facilitating the activity of midfrontal cortex or the connected network, results in better performance in interpersonal motor interactions (Boukarras et al., 2022; Sacheli et al., 2023). Neurophysiological evidence highlights the crucial role of the anterior cingulate cortex in generating midfrontal theta and error-related negativity (ERN) during performance monitoring (Cavanagh & Frank, 2014; Jocham & Ullsperger, 2009). The early Pe, instead, is suggested to originate from the anterior regions of the ACC (Holroyd & Coles, 2002), and has been associated to attentional reorientation (Ridderinkhof et al., 2009), while the late Pe may originate from the insular cortices, signaling error awareness (Klein et al., 2013).

Among these electrocortical markers associated with the activity of the performance monitoring system, ERN and midfrontal theta oscillations are proposed to be influenced by dopaminergic activity (Parker et al., 2015). Notably, studies have demonstrated that the midfrontal cortex receives dense dopaminergic projections from the ventral tegmental area (VTA), (Ullsperger et al., 2014). However, the impact of dopamine on modulating interpersonal performance monitoring system’s activity through neuronal synchronization remains understudied. To address this gap, examining patients with PD under various pharmacological conditions, such as during dopaminergic medication (PD ON) and after drug withdrawal (PD OFF), allows for a direct exploration of the dopaminergic system’s contribution to interpersonal performance monitoring. A recent study revealed that patients with PD in ON exhibited the expected increase in midfrontal theta power following the observation of erroneous actions, while patients with PD in OFF did not. These findings imply that the depletion of dopamine has a significant impact on this neurophysiological marker associated with performance monitoring (Pezzetta et al., 2023).

The role of dopamine in modulating the performance monitoring system’s activity during interpersonal motor interactions remains less explored. One behavioural study (Era et al., 2023) has shown that patients with PD in OFF manifest difficulties in coordinating their actions with a virtual partner, in conditions requiring continuous monitoring adaptation to its actions. Interestingly, this ability is maintained when patients take dopaminergic medications. Nevertheless, the connection between these observed difficulties and potential alterations in the neurophysiological markers associated with interpersonal performance monitoring remains uninvestigated.

In addressing this research gap, the current investigation involved the examination of patients with PD both in the ON and OFF condition, alongside a group of healthy controls (HCs). Electroencephalography (EEG) was recorded while participants were tasked with grasping a bottle-shaped object in synchrony with a virtual partner (Fig. 1). This task included two distinct conditions: (i) a Cued condition, where participants had advance knowledge of their designated grasp location, and (ii) an Interactive condition, requiring participants to coordinate their action according to the virtual partner’s movement by either imitating or complementing its movement (thus demanding continuous monitoring adaptation to the virtual partner’s actions). Moreover, we included a correction factor, involving sudden changes in the virtual partner’s actions (VP’s correction), prompting participants to flexibly adapt to the movements of their virtual partner in the Interactive condition. Our hypotheses build upon prior research and anticipates replicating findings that suggest patients with PD in OFF encounter challenges in coordinating with a virtual partner, particularly in the Interactive compared to the Cued condition (Era et al., 2023). Furthermore, we hypothesize that these coordination difficulties may be paralleled by changes in electrocortical markers associated with interpersonal performance monitoring and adaptation, particularly midfrontal theta oscillations, which have been previously shown to be modulated by dopaminergic contributions (Pezzetta et al., 2023). We complement classical univariate analyses (focusing on electrocortical markers of interpersonal performance monitoring, such as the ERN, Pe, and midfrontal Delta-Theta power, as described by Moreau et al., 2020) with multivariate pattern analysis (MVPA) to describe whether different interactive conditions are characterized by different cortical processing patterns as a function of dopaminergic impairments. Specifically, we aim to: i) determine if and when a classifier can distinguish between the Interactive and Cued conditions across the three groups based on whole-brain EEG data; ii) assess whether classifier accuracy varies between PD ON, PD OFF, and healthy controls (HCs); and iii) evaluate whether the classifier can differentiate between the Interactive and Cued conditions before the VP’s Correction, especially in the PD OFF condition, suggesting a modulation of proactive cognitive control (i.e. the anticipation of cognitively demanding events, such as the possible VP’s Correction, before they occur) in patients with PD.

**Figure 1.**
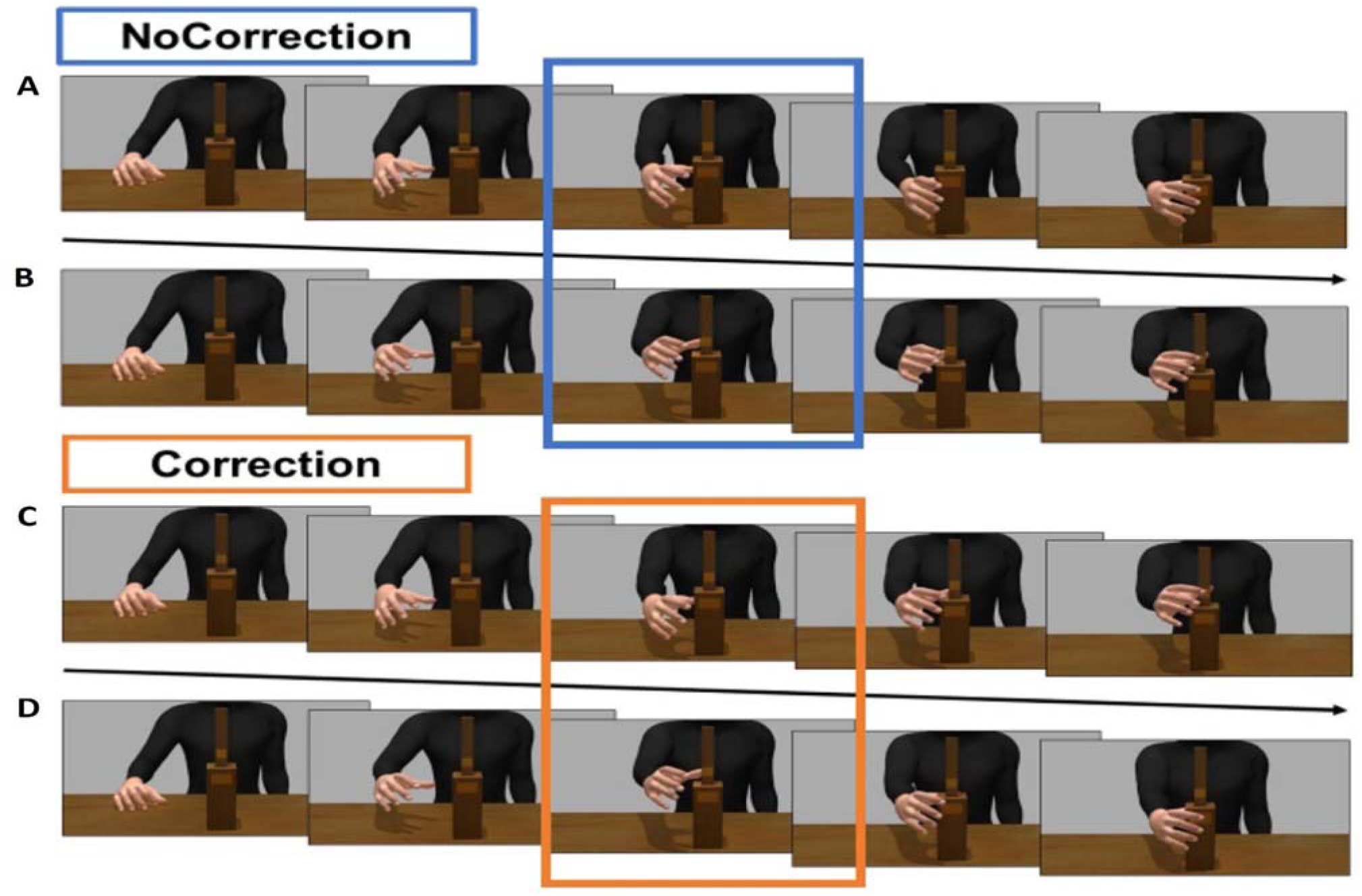
Examples of the sequence of frames for each type of VP’s movement: A) Power Grasp; B) Precision Grasp; C) Correction Power to Precision Grasp and D) Correction Precision to Power Grasp. The middle frame of each sequence represents the time when the EEG trigger to lock the Correction (orange) or NoCorrection (blue) to was sent.

Results showed that patients with PD in OFF exhibited worse behavioral performance in the Interactive condition compared to patients with PD in ON and to HCs. Univariate electroencephalography analyses revealed higher midfrontal Delta-Theta activity in the Interactive condition in PD OFF compared to PD ON. Additionally, in all groups, better behavioral performance in Interactive trials was associated with higher midfrontal Delta-Theta synchronization, possibly indicating an attempt towards compensation for disrupted performance monitoring abilities in PD OFF. Finally, the multivariate classification analysis demonstrated that a classifier could distinguish between Interactive and Cued trials based on whole-brain EEG patterns before the VP’s correction, with superior performance observed in the PD OFF condition. These findings: i) underscore the distinct involvement of neural networks involved in interpersonal performance monitoring during Interactive compared to Cued trials; ii) highlight the modulation of neural correlates associated with interpersonal performance monitoring by the dopaminergic system; and iii) support the notion that proactive cognitive control is impaired in patients with Parkinson’s disease (PD) (Kricheldorff et al., 2023).

## RESULTS

### Behavioural performance in interpersonal motor interactions is modulated by dopamine depletion

To investigate the role of the dopaminergic system during interpersonal motor interactions, we analysed the performance of patients with PD both in the ON and OFF condition, as well as the performance of a group of HCs, in a highly ecological and well-validated Joint-Grasping task. We used Grasping Asynchrony, i.e. the absolute time delay between the participant’s and the virtual partner’s touch time on a bottle-shaped object (Fig. 1), to measure the success of the interaction. This dependent measure was corrected by subtracting the performance in a control condition (i.e., Cued condition), which does not rely on interpersonal performance monitoring, from the performance in the Interactive condition.

Behavioural (i.e., Grasping Asynchrony) results are plotted in Fig. 2, with the main difference between groups estimated in terms of Grasping Asynchrony. In detail, the 2 Condition (PD ON/OFF) × 2 Interaction type (Complementary/Imitative) × 2 Movement Type (Precision/Power grip) × 2 Correction (Correction/NoCorrection), within-participants ANOVA on Grasping Asynchrony (i.e., Interactive minus Cued Condition) showed a significant main effect of the Condition factor [F(1, 14) = 5.876, p = .029, ηp² = .296], with patients performing significantly worse in the motor interaction task when they were in the OFF (M = 313.19 ms, SD = 216.35) than in the ON medication condition (M = 239.53 ms, SD = 192.91).

**Figure 2.**
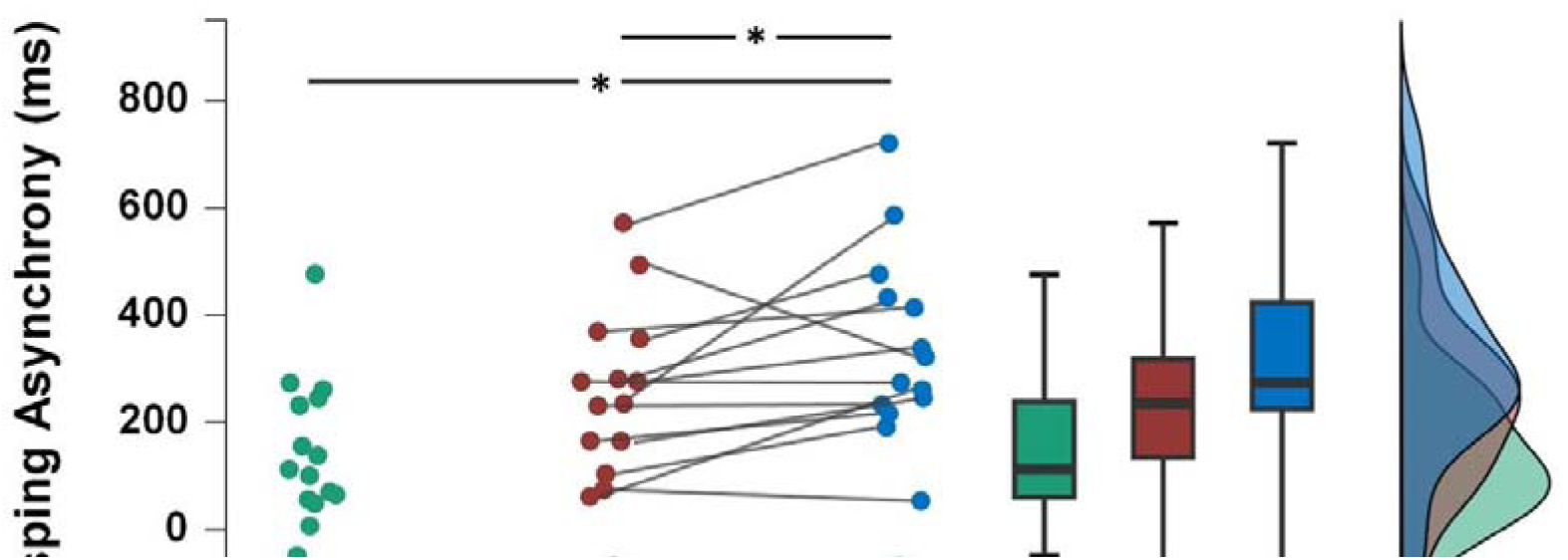
Raincloud plot displaying participants’ performance in the motor interaction task. PD OFF patients performed significantly worse compared to themselves in the ON condition (p. = .029), as well as to HCs (p. = .01). No significant difference was found between HCs and PD ON patients.

In the 2 Group (PD ON/HC) x 2 Interaction type (Complementary/Imitative) × 2 Movement Type (Precision/Power grip) × 2 Correction (Correction/NoCorrection) mixed ANOVA on Grasping Asynchrony (Interactive minus Cued Condition) the Group factor did not reach statistical significance (p = .103).

The 2 Group (PD OFF/HC) x 2 Interaction type (Complementary/Imitative) × 2 Movement Type (Precision/Power grip) × 2 Correction (Correction/NoCorrection) mixed ANOVA on Grasping Asynchrony (Interactive minus Cued Condition) showed a significant main effect of the Group factor [F(1, 28) = 7.384, p = .011, ηp² = .209], showing that patients in the OFF medication condition performed significantly worse (M = 313.19 ms, SD = 216.35) than HCs (M = 146.47 ms, SD = 172.00). The Group factor did not interact with any within-subject factor.

### Oscillatory dynamics, but not ERPs, associated with interpersonal performance monitoring significantly increase in patients with PD during OFF medication

At the neurophysiological level, we measured different electrocortical markers of interpersonal performance monitoring, such as the ERN and the Pe (in the time-domain), and the midfrontal Delta-Theta power (in the time-frequency domain), as described by Moreau and colleagues (2020).

Starting from the analysis on the amplitude of the Pe component, the 2 Condition (PD ON/OFF) x 2 Interactivity (Interactive/Cued) x 2 Correction (Correction/NoCorrection), within-participants ANOVA revealed a significant interaction between the three main factors (Condition x Correction x Interactivity) [F(1, 14) = 6.077, p = =.027, ηp² = .303]. Post-hoc tests indicated that the Pe amplitude was significantly larger in the ON condition patients during Interactive-Correction trials than during both Interactive-NoCorrection trials (p < .001) and Cued-Correction trials (p < .001), as well as for OFF condition patients during Interactive-Correction trials compared to Interactive-NoCorrection (p < .001) and Cued-Correction ones (p < .001). However, no significant differences emerged between the OFF and ON condition.

The 2 Group (PD ON/HC) x 2 Interactivity (Interactive/Cued) x 2 Correction (Correction/NoCorrection) mixed ANOVA on Pe amplitudes revealed a significant interaction between the three main factors (Group x Correction x Interactivity) [F(1) = 4.691, p = .039, ηp² = .143]. Post-hoc tests indicated that the Pe amplitude was significantly larger in the ON condition patients during Interactive-Correction trials than during Interactive-NoCorrection trials (p < .001) and Cued-Correction trials (p < .001), as well as in the HCs during Interactive-Correction trials compared to Interactive-NoCorrection (p < .001) and Cued-Correction ones (p < .001). However, no significant differences emerged between the ON and HCs groups.

The 2 Group (PD OFF/HC) x 2 Interactivity (Interactive/Cued) x 2 Correction (Correction/NoCorrection) mixed ANOVA on Pe amplitudes did not reveal any significant main effect or interactions with the factor Group, all ps > 0.56. All time-domain results are plotted in Figure 3.

**Figure 3.**
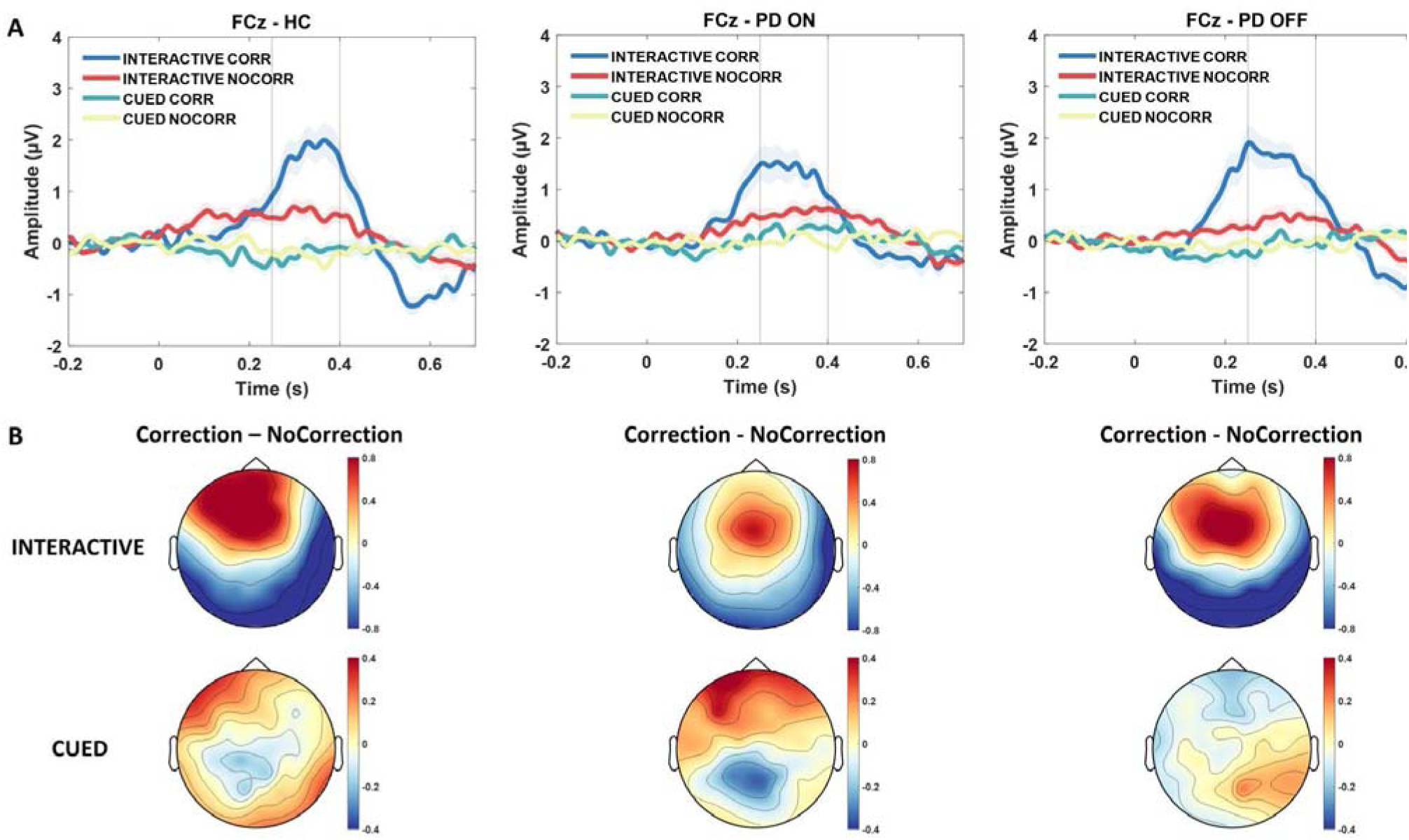
A) Grand averages of the Pe component over FCz in all experimental conditions, divided by group (HC, PD ON/OFF). Data are time-locked to the correction of the Virtual Partner equivalent frame when no correction occurred). B) Topographies of the difference between component (from 250 to 400 ms after the VP’s correction) in the Correction compared (or the the Pe to the NoCorrection conditions, divided between Interactive and Cued conditions as well as between groups.

Focusing on the midfrontal oscillatory activity in the Delta-Theta band (2-7 Hz) typically related to action monitoring, the 2 Condition (PD ON/OFF) x 2 Interactivity (Interactive/Cued) x 2 Correction (Correction/NoCorrection) within-participants ANOVA showed a significant interaction between the Condition and Interactivity factors [F(1, 14) = 7.553, p = .016, ηp² = .350]. Post-hoc tests indicated that Delta-Theta synchronization was larger during Interactive trials compared to Cued one in both ON and OFF condition patients. In details, Delta-Theta synchronization was larger for ON condition patients in the Interactive trials compared to the Cued ones (p = .004), as well as for OFF condition patients in the Interactive trials compared to ON condition patients in the Interactive trials (p = .048) and also compared to Cued ones (p. < 001). This pattern of results suggests that OFF compared to ON condition patients show a Delta-Theta increase in the Interactive condition.

The 2 Group (PD ON/HC) x 2 Interactivity (Interactive/Cued) x 2 Correction (Correction/NoCorrection) mixed ANOVA did not show any significant main effect or interaction of the Group factor, all p > 0.21.

The 2 Group (PD OFF/HC) x 2 Interactivity (Interactive/Cued) x 2 Correction (Correction/NoCorrection) mixed ANOVA showed a significant interaction between the Group and Correction factor [F(1) = 5.128, p = .031, ηp² = .155]. Post-hoc tests indicated that Delta-Theta synchronization was larger during Correction trials compared to NoCorrection ones in both OFF and HCs. In detail, Delta-Theta synchronization was larger for OFF condition patients during Correction trials compared to NoCorrection ones (p = .005), as well as for HCs during Correction trials compared to NoCorrection ones (p < .001). However, no significant differences emerged between the OFF and HCs groups. This pattern indicates that for both OFF condition and HCs, both in Interactive and Cued trials, dealing with a Correction increased midfrontal Delta-Theta synchronization.

Lastly, confirming the pattern found in the OFF/ON analysis, also the interaction between the Group and Interactivity factors resulted significant [F(1) = 4.797, p = .037, ηp² = .146]. Post-hoc tests indicated that Delta-Theta synchronization was larger during Interactive trials compared to Cued one in both OFF and HCs. In detail, Delta-Theta synchronization was larger for OFF condition patients during Interactive trials compared to Cued ones (p < .001), as well as for HCs during Interactive trials compared to Cued ones (p = .002). However, no significant differences emerged between the OFF and HCs groups.

As control analyses, we also run the same set of statistical comparisons on ERD/ERS values in the Alpha (8-13 Hz) and Beta (14-30 Hz) band. None of these analyses relieved significant main e fects or interactions. Figure 4 represents the ERD/ERS for each group (HC, PD ON and OFF) in the Interactive and in the Cued block.

**Figure 4.**
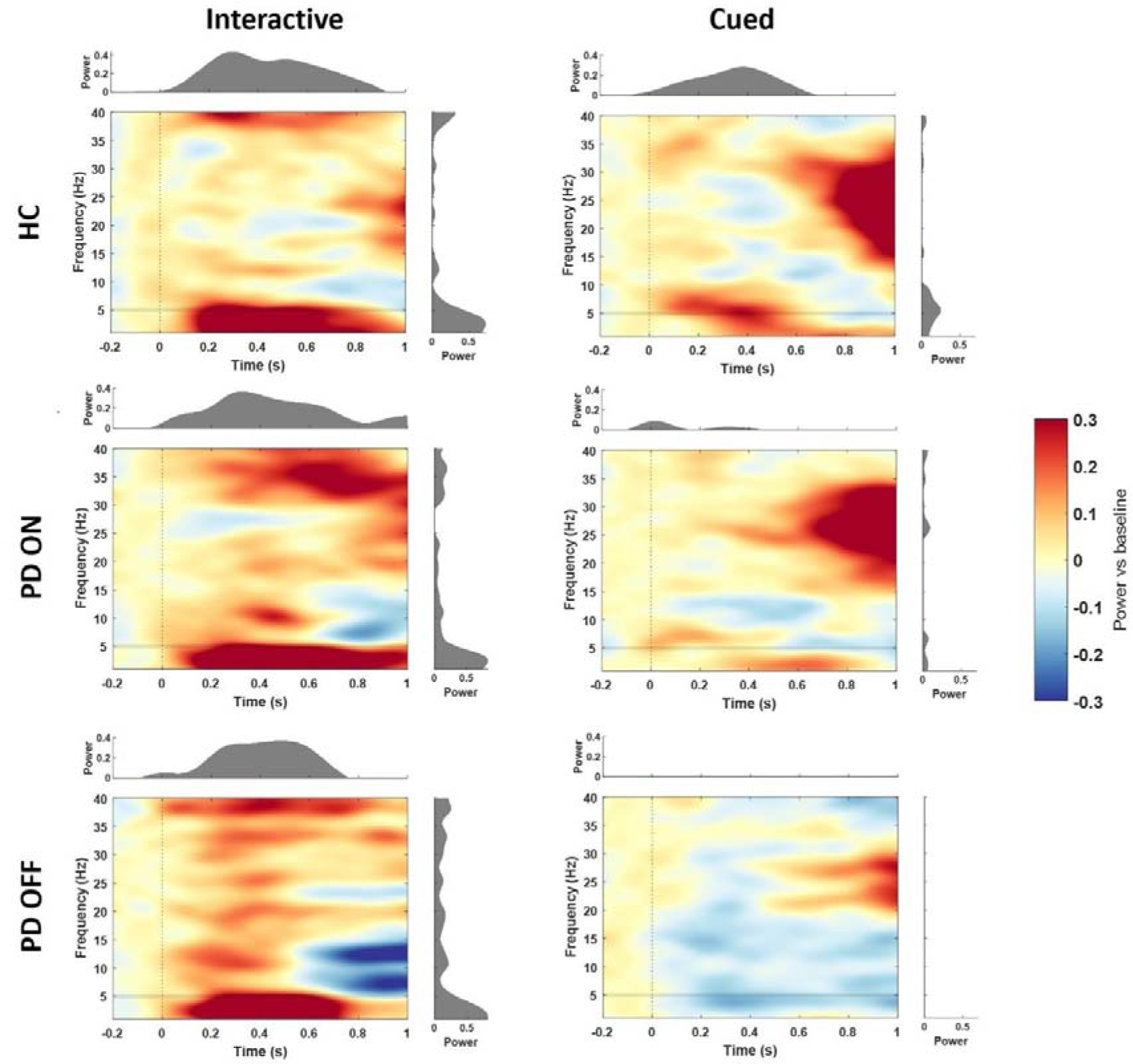
Power change relative to baseline over electrode FCz for Interactive and Cued conditions, separately for each group. Left columns: time-frequency representations for all the Interactive conditions (collapsing Correction and NoCorrection trials). Right columns: time-frequency representations for all the Cued conditions with no correction (collapsing Correction and NoCorrection trials).

We also run the difference in power, for each group, between Interactive and Cued trials. We ran three separate ANOVAs comparing such differences in power between PD ON and OFF (with GROUP as within-subject factor), and between PD ON and HCs, and OFF and HCs (with GROUP as between-subject factor). These analyses show that the difference in Delta-Theta power between Interactive and Cued trials is significantly stronger in PD OFF patients compared to themselves in the ON condition (p = .008) as well as compared to the data from the HCs (p = .043). See Figure 5 for a representation of the difference in power between Interactive and Cued trials for each group (for the respective topographies, see Fig. S1/2/3).

**Figure 5.**
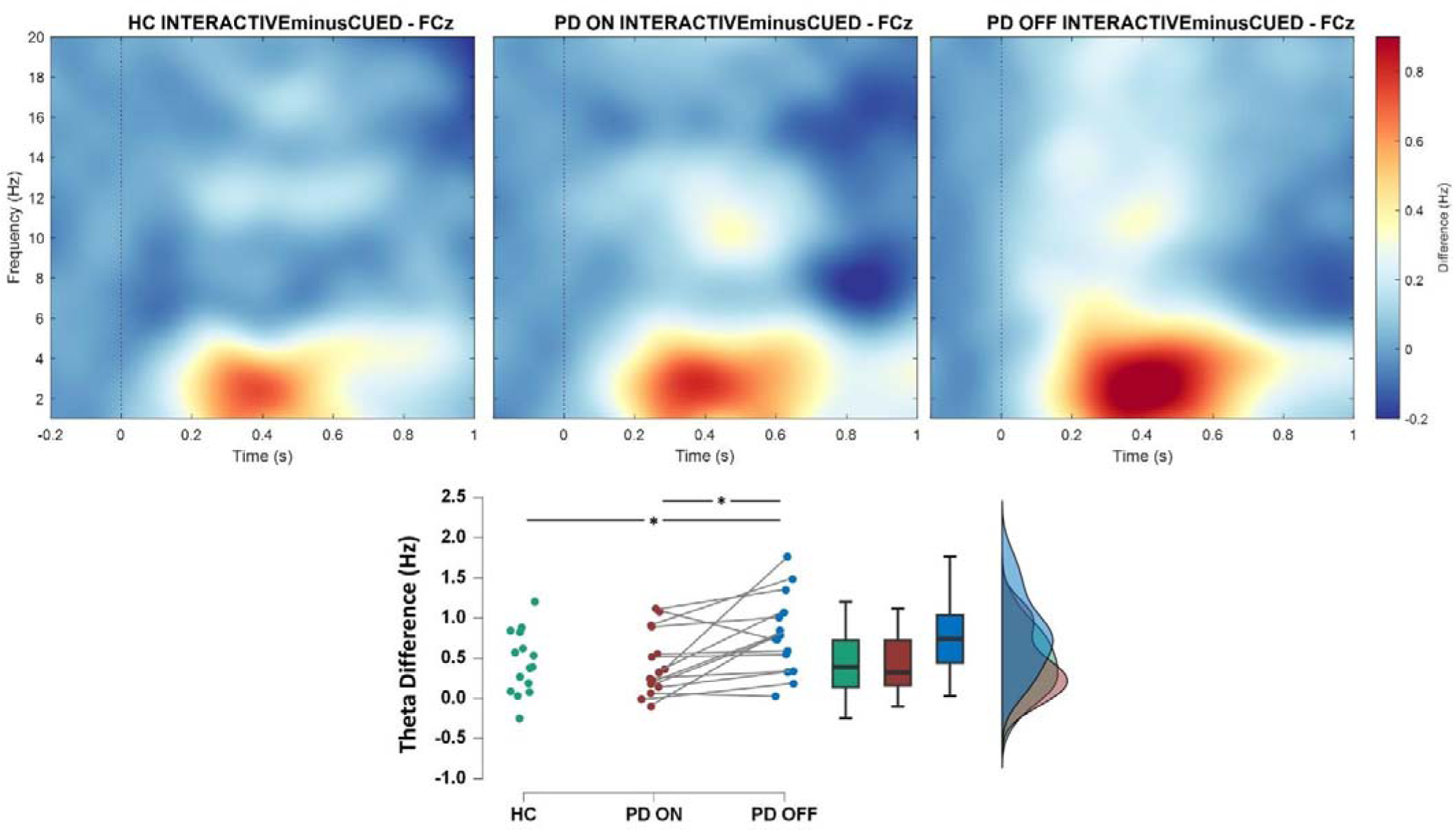
Time-frequency representations showing a main effect of the group factor. PD OFF patients showed a stronger difference between the frontocentral synchronization in Delta-Theta, when subtracting the Cued from the Interactive condition, compared to themselves in the ON condition, or to the HCs group.

### Midfrontal Delta-Theta ERS is associated with behavioral performance in the Interactive condition

To test the influence of participants’ midfrontal Delta-Theta synchronization on the ability to perform the Joint-Grasping task, we entered single-trial data of midfrontal Delta-Theta ERS as continuous predictor in a linear mixed model, while single-trial Grasping Asynchrony scores represented the dependent variable. The model also included as categorical predictors the Condition (Interactive, Cued) and Group (PD ON/OFF, HC). Only significant main effects or interactions with the continuous predictor Delta-Theta synchronization are reported here. For all the other main effects and interactions see Table S16. The model showed a significant interaction between Condition and Delta-Theta ERS (F(1,9492.8)=8.32 p=.004). Simple slope analysis showed that only the slope of the Interactive condition was significantly different from zero as a function of Delta-Theta synchronization [LCI −4.56 - UCI −0.62]. More specifically the higher midfrontal Delta-Theta ERS the lower the Grasping Asynchrony in the Interactive condition (that is the better participants’ performance) (Figure 6).

**Figure 6.**
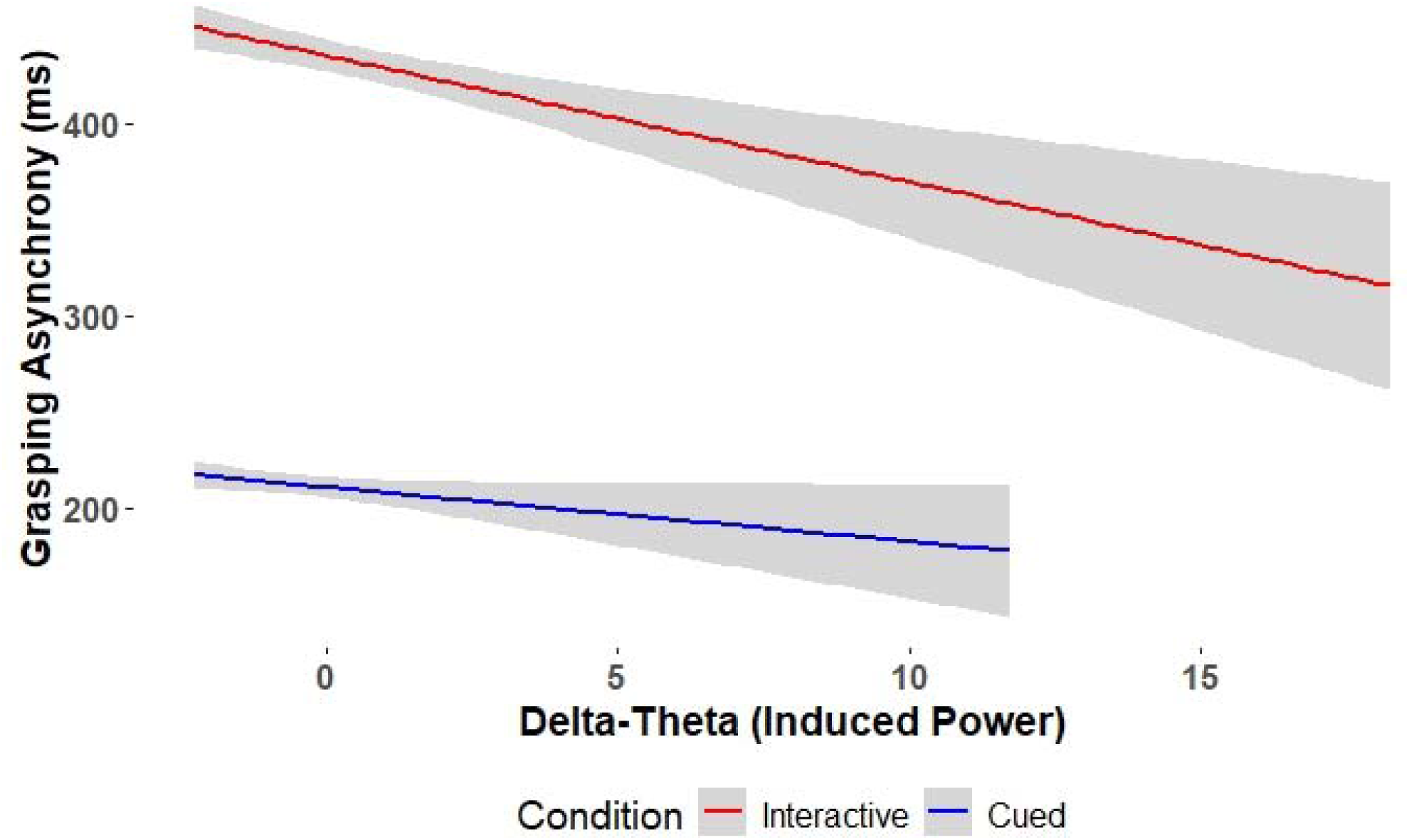
Results of the analysis on the interaction between midfrontal Delta-Theta ERS and behavioural performance in each condition (Interactive vs Cued). The higher the midfrontal Delta-Theta synchronization, the better participants’ behavioural performance (i.e., the lower the Grasping Asynchrony) in the Interactive condition.

### EEG decoding highlights differences in the neural patterns related to early stages of interpersonal monitoring dependent on dopamine depletion

A Multivariate Pattern Analysis (MVPA) was conducted to determine whether and when neurophysiological indices associated with the Interactive condition could be distinguished from those linked to the Cued condition across different groups. This analysis aimed to investigate the role of dopamine in differentially influencing neural processing related to performance monitoring based on varying levels of task interactivity. This corresponds to assessing if and how a classifier (i.e., a LDA model) would be able to distinguish the single-trial EEG patterns related to the monitoring of the VP during the Interactive compared to the Cued task, in each experimental group. The results of these analyses are shown in Figure 7. In the lower panel, the results from the classification are shown separately for each experimental group, highlighting that Interactive and Cued trials could be decoded significantly better than chance for each group. Specifically, for all groups, the LDA performance was significantly different than chance level as early as ∼1500 ms before the VP correction, both when decoding Interactive and Cued trials (significant clusters time windows: HC Interactive [-1.61 to 1s], Cued [-1.56 to 1s]; PD ON Interactive [-1.45 to 1s], Cued [-0.84 to 1s]; PD OFF Interactive [-1.59 to 1s], Cued [-1.78 to 1s] – as shown by the bold dotted lines at the top and bottom of the decoding curves in Fig. 7 panel B). The classification pattern remained sustained over time and reached its maximum peak after 0 ms (i.e. when the correction occurs).

**Figure 7.**
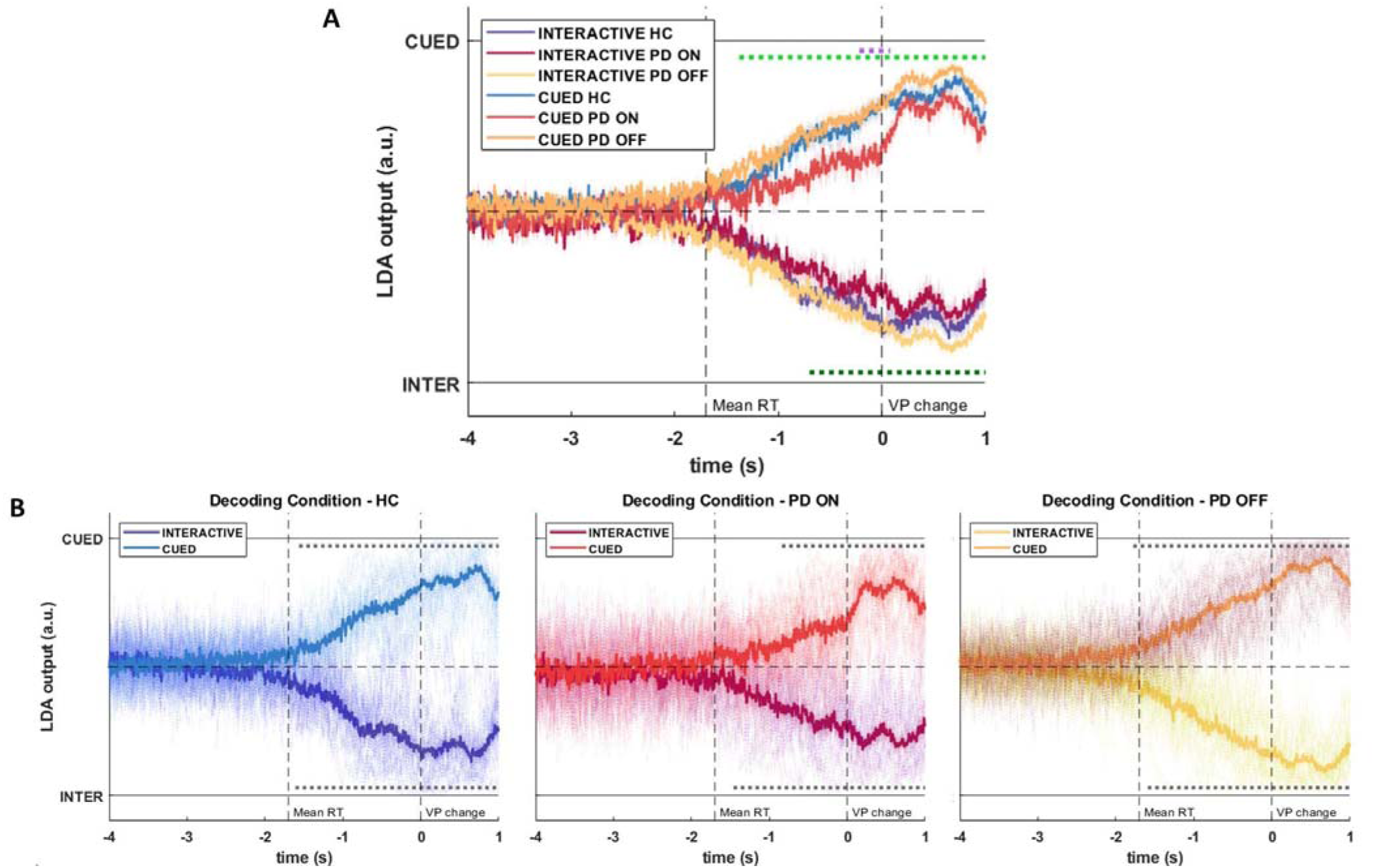
A) MVPA results highlighting the performance of the LDA classifier (i.e., classifier’s output) in distinguishing trials in the Interactive compared to the Cued condition, for each experimental group. Dotted lines at the top indicate time-windows when the decoding performance for the Cued condition significantly differed between PD OFF and PD ON (light green line), and between PD OFF and HCs (purple line). The dotted line at the bottom indicates the time-window when the decoding performance for the Interactive condition significantly differed between PD OFF and PD ON (dark green l ne). B) Decoding performance (i.e., classifier’s output) plotted separately for each group. Dotted lines at the top and bottom highlight time-windows with significantly (p. < .05, cluster-corrected) better than chance classification. “Mean RT” = average subject’s movement start.

In order to compare the performance of the classifier across groups, i.e., the effectiveness of the linear combination of features that allow the classification, we performed, separately for Interactive and Cued trials decoding, 1) a dependent samples t-test across time between PD ON and PD OFF, and 2) two independent samples t-tests across time between PD ON and HCs, and PD OFF and HCs with statistical significance being assessed through non-parametric cluster-based permutations (n = 1000; Maris & Oostenveld, 2007). We found that the performance of the classifier in decoding Cued trials was significantly better when patients were OFF compared to ON medication, in a 2.4 seconds time window between −1.38 and 1 s around the correction of the VP (light green dotted line on top of Fig. 8, panel A), as well as compared to the performance for the HCs group, in a much shorter .30 seconds time window between −0.22 and 0.08 s around 0 (purple dotted line on top of Fig. 7, panel A). Moreover, with regard to Interactive trials, the performance of the classifier was significantly better when patients were OFF compared to ON medication, in a 1.7 seconds time window between −0.70 and 1 s around VP’s correction (dark green dotted line at the bottom of Fig. 7, panel A).

**Figure 8.**
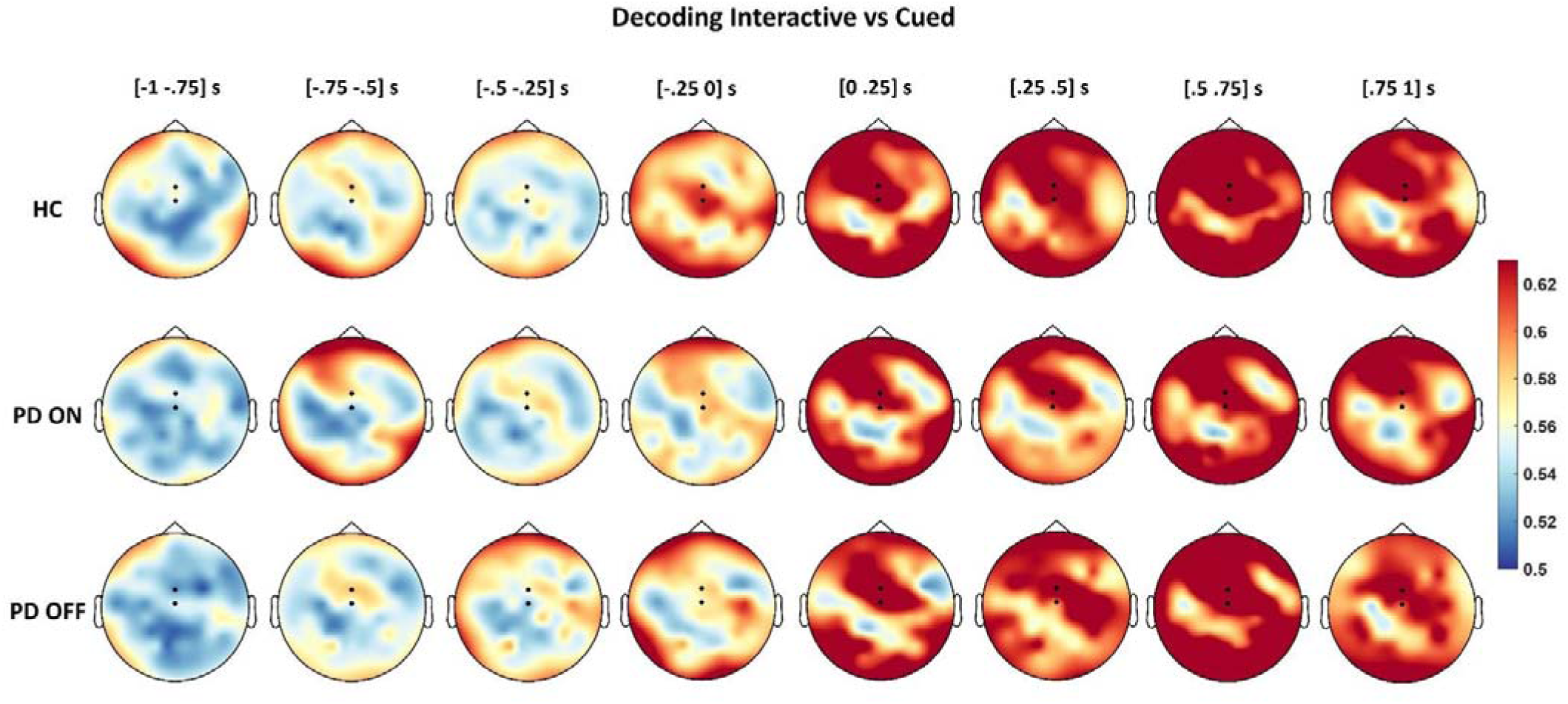
Topographies of the searchlight analysis results showing the temporal and spatial distribution of accuracy values for the binary classification of Interactive vs Cued trials separately for each group (HC/PD ON/ PD OFF). In all groups, fronto-central sites consistently contributed to the classifier’s performance from ∼ 500 ms before the VP’ correction (time 0). The two dots on each topography represent electrodes Cz and FCz.

In order to test the weight of different electrodes contributing to classification performance, we ran a searchlight analysis across time and electrodes with the same parameters implemented for the classification across time points (i.e., LDA classifier, k-fold cross-validation), this time quantifying the classifier’s performance in terms of % accuracy. These results are plotted in Figure 8 as topographies in eight time-windows of interest (from −1 to 1 second around the timepoint of the correction, i.e., 0, in steps of 250 ms), showing that the classifier relied on a distributed cluster of electrodes to decode Interactive vs Cued trials, with a fronto-central cluster being most prominent from ∼500 ms before the VP’s correction (or the equivalent frame in NoCorrection trials) in all groups. This approach complements the results from the univariate analysis and localizes the results from the binary classification over time (Fig. 8), providing further evidence for a modulation of the performance monitoring system by the dopaminergic system.

## DISCUSSION

To explore the impact of dopamine on behavioural and electrocortical signatures of interpersonal performance monitoring, we conducted an experiment involving patients with Parkinson’s Disease (PD) in two conditions, i.e., while regularly receiving dopaminergic medication (PD ON), and after the withdrawal from medication (PD OFF). Participants were tasked with coordinating their actions with a virtual partner in conditions where continuous monitoring of and adaptation to its actions are (Interactive condition) or are not (Cued condition) required.

### Dopamine depletion worsens behavioural performance during interpersonal motor interactions

Behavioural results showed that, after controlling for patients’ motor and cognitive ability to perform the task in Cued (control) trials, PD OFF performed worse in the Interactive trials compared to themselves in the ON condition. In keeping with Era et al. (2023), this confirms the role of the dopaminergic system in supporting the ability to adapt one’s movements to those of an interaction partner, particularly in situations necessitating continuous monitoring of its actions. Indeed, PD OFF exhibited interpersonal coordination difficulty in contexts requiring flexibility in adapting to others’ movements. In the Cued condition, instead, participants were already instructed on which action to perform and likely dedicated fewer cognitive resources to monitoring the partner’s behaviour. Furthermore, PD OFF performed less effectively than healthy controls (HCs), while in the ON condition they did not differ from HCs in the Interactive trials. This result corroborates the notion that dopaminergic medication facilitates the ability of patients with PD to successfully coordinate with a virtual partner when predictions and monitoring of its actions are required (Era et al., 2023).

### Univariate analyses unveil dopamine-modulated electrocortical markers of interpersonal performance monitoring in the time-frequency domain

Concerning the electrocortical markers associated with interpersonal performance monitoring, results revealed that only midfrontal Delta-Theta activity was influenced by dopamine levels. Indeed, higher midfrontal Delta-Theta activity was observed in PD OFF compared to PD ON, aligning with the proposed role of dopamine in performance monitoring and the regulation of social predictive processes (Solié et al., 2022). Interestingly, however, PD OFF showed an hyperactivation of the performance monitoring system, highlighted by increased synchronization in the frontal Delta-Theta band during the Interactive trials, compared to PD ON patients. This is in contrast with what previously found in studies investigating the role of dopamine in modulating the activity of the performance monitoring system in individual/observational tasks, showing a reduced activity of the performance monitoring system in PD OFF (Pezzetta et al., 2023). Moreover, when looking at midfrontal Delta-Theta activity in the Interactive minus the Cued condition, higher midfrontal Delta-Theta activity was observed in PD OFF compared to PD ON as well as compared to HCs. Although not statistically significant in univariate analysis, the pattern of results in the Cued condition mirrored those found in prior studies (Pezzetta et al., 2023, a study in which patients with PD had simply to observe a movement after a cue but did not have to actively adjust their own movement in order to anticipate other’s actions), showing decreased synchronization in the frontal Delta-Theta band in PD OFF compared to themselves in the ON condition and HCs. Thus, dopamine may influence the activity of the performance monitoring system differently depending on the context’s interactivity level. Specifically, the performance monitoring system is more active in PD OFF when there is a greater need to monitor and adapt to others’ actions (Interactive condition) compared to situations where such monitoring and adaptation are less necessary (Cued condition) or when passively observing a movement performed by a virtual arm from a first-person perspective (Pezzetta et al., 2023).

### Higher Delta-Theta synchronization is associated with better performance in the Interactive condition, suggesting its potential compensation function in PD OFF

While the Interactive condition was associated with overall worse behavioral performance in PD OFF. When directly examining the relationship between single-trial midfrontal Delta-Theta synchronization and patients’ ability to perform the Joint-Grasping Task, in all groups, he higher the midfrontal Delta-Theta synchronization the better the behavioral performance (i.e., lower Grasping Asynchrony). Functional Magnetic Resonance Imaging findings indicate that behavioural adjustments trigger activations in both the anterior cingulate cortex (ACC) and the lateral prefrontal cortex (LPFC) (Kerns et al., 2004). The ACC is thought to oversee individuals’ actions and, when needed, communicate with the LPFC to engage cognitive control mechanisms to improve subsequent performance (Kerns et al., 2004). Crucially, theta oscillations are implicated in facilitating neural communication between midfrontal and frontal areas during motor adaptation (Oehm et al., 2014). Reduced functional connectivity between the ACC and LPFC has been shown in patients with PD, in tasks tapping on performance monitoring abilities (de Bondt et al., 2016). This reduced connectivity may explain the poorer behavioral performance of patients with PD (especially in the OFF condition) in the Interactive condition of the present study, which requires continuous monitoring and adaptation to others’ behavior. Conversely, higher midfrontal Delta-Theta synchronization in PD OFF patients may represent an attempt to compensate for disrupted communication between the ACC, involved in monitoring the partner’s actions during interpersonal motor interactions, and the frontal regions involved in behavioral adaptation.

### Dopamine depletion does not modulate ERPs associated with interpersonal performance monitoring

The present results of the Correction trials also revealed a detectable Pe recorded over fronto-central electrodes in all groups. Notably, the lack of a significant difference in the Pe response among groups aligns with the suggestion of an independent generation of this component from the dopaminergic system (Falkestein et al., 2001). This evidence aligns with a previous study indicating that dopamine depletion in patients with PD is associated with a selective modulation of midfrontal theta activity in tasks relying on performance monitoring, while leaving the Pe component unaffected (Pezzetta et al., 2023). Conversely, we did not detect any late Pe over the parietal electrodes, which aligns with previous studies suggesting that in older adults, a component shift from posterior to anterior areas may occur (Overbeek et al., 2005; Pezzetta et al., 2023). Surprisingly, we did not detect any ERN, an error-related marker present in previous studies investigating interpersonal performance monitoring in young adults (Moreau et al., 2020; 2023). However, the ERN was absent also in previous studies investigating older populations with tasks tapping into the activity of the performance monitoring system (Spinelli et al., 2022; Pezzetta et al., 2023).

### EEG decoding analyses further support dopamine role in differentiating the neural patterns related to interpersonal monitoring before the VP’s Correction

Finally, we complemented our univariate analyses with a multivariate approach, showing that a classifier (i.e., a LDA model) was able to distinguish between Interactive and Cued trials based on the EEG patterns in all three groups, as early as 1500 ms before the VP’s Correction or the corresponding time window in No-Correction trials. Moreover, the performance of the classifier was significantly better when relying on the neural patterns recorded in patients with PD in OFF compared to themselves in the ON condition, in both the Cued and Interactive conditions (Fig. 7, panel A). Thus, the absence of medication leads to a significantly stronger differentiation of the neural correlates distinguishing interpersonal interactions requiring continuous monitoring and adaptation to others’ actions (i.e., Interactive) to conditions not tapping on these abilities (i.e., Cued). It is noteworthy that the classifier effectively distinguished between the Interactive and Cued conditions well before the VP’s Co rection, with better performance observed in PD OFF compared to PD ON. This, coupled with the finding that patients with PD, particularly in the OFF condition, showed altered behavioral performance and EEG activity in the Interactive condition regardless of whether the virtual partner performed a movement Correction, supports the notion that proactive cognitive control is impaired in patients with PD (Kricheldorff et al., 2023). In fact, proactive cognitive control, which involves anticipat ng and preventing cognitively demanding events (the possible VP’s Correction in this case) before they occur, is a common factor in both Interactive-Correction and Interactive-NoCorrection trials. Finally, the classifier’s better performance in the PD OFF condition compared to the ON one for both Interactive and Cued trials further support the idea that dopamine influences neural activity across various levels of interactivity.

Interestingly, the results of the searchlight classification highlighted the involvement of fronto-central sites in distinguishing Interactive and Cued trials. Notably, this complements the analyses of midfrontal Delta-Theta synchronization, where we focused on induced, non-phase locked, power (thus, without looking at the evoked, phase-locked, patterns used by the classifier). All in all, the MVPA results: i) further support the different involvement of the neural network involved in interpersonal performance monitoring in the Interactive compared to the Cued trials; ii) most importantly, they highlight the modulation of the neural correlates of interpersonal performance monitoring by the dopaminergic system; iii) by demonstrating that the classifier effectively distinguished between the Interactive and Cued conditions well before the VP’s Correction, with better performance in the PD OFF condition, they provide evidence that proactive cognitive control is impaired in patients with PD.

In conclusion, our study underscores the crucial role of the dopaminergic system in influencing both behavioral and electrophysiological responses during interpersonal performance monitoring. The findings identify electrocortical markers modulated by dopamine, associated with performance monitoring in interactive tasks. These insights may guide the development of clinical interventions combining non-invasive brain stimulation with interpersonal motor interactions, potentially enhancing neurophysiological markers of interpersonal performance monitoring and improving motor interaction abilities in patients with PD.

## METHODS

### Participants

Sixteen patients affected by Parkinson’s Disease (i.e., PD) were involved in the study. The sample size was determined through a prospective power analysis performed with the software More Power (Campbell and Thompson, 2012). We inserted as expected effect size the partial eta squared value (0.38) observed in (Pezzetta et al., 2023) where the electrocortical signatures associated to the activity of the performance monitoring system with the activity of the performance monitoring system were studied in patients with PD. The analysis indicated that a 2 × 2 × 2 within-between design, a power of 0.80 and a partial eta squared value of 0.38 (as is computed from Pezzetta et al., 2023), required a sample size of 16 participants.

All participants had normal or corrected-to-normal vision and were naive as to the purpose of the experiment. Patients with PD were recruited for the study according to the following inclusion criteria: (i) diagnosis of idiopathic PD (United Kingdom Parkinson’s Disease Society brain bank criteria, UPDRS) (ii) absence of dementia (Mini Mental State Examination, MMSE above or equal to 25); (iii) absence of other neurological and psychiatric diseases; (iv) assumption of taking daily doses of dopamine or a dopamine agonist (L-Dopa equivalent doses are reported in Table 1). One patient was excluded from the final sample, due to excessive muscular artifacts in the EEG, leading to the exclusion of >25% of the ICA components. The final sample included 15 patients (14 males, 1 female, group average age = 67.47 ± 7.48 years; group average years of education = 14 ± 3.87; group average MMSE = 29.4 ± 0.63). Besides being based on a prospective power analyses, the sample size is the same as in previous studies investigating performance monitoring in patients with PD (Pezzetta et al., 2023, n = 15, Era et al., 2023, n = 15). The socio-demographic and clinical characteristics of patients who participated in the study are reported in Table 1. Moreover, 15 HCs were involved in the study as a control group (13 males, 2 females, group average age = 67.2 ± 5.75 years; group average years of education = 14.67 ± 3.29; group average MMSE = 28.9 ± 0.9). Age, years of education and MMSE scores did not differ between the PD and HCs groups (p = 0.92, p = 0.64 and p = 0.57, respectively).

**Table 1.** Summary of demographics and clinical scores of the patients with PD. Age: age in years, Education: education in years. Mean values of the UPDRS-III motor scale and H&Y scale for disease progression of patients with PD tested on- (with dopaminergic medication) and off-condition (without dopaminergic medication). MMSE mini-mental state examination, MMPSE mini-mental Parkinson state examination, L-Dopa equivalents: daily dose of levodopa and/or dopamine agonists. NA corresponds to missing data.

| N | Gender | Age | Education | UPDRS_ON | UPDRS_OFF | H&Y_ON | H&Y_OFF | MMS E | MMPSE | L-Dopa equivalents |
| --- | --- | --- | --- | --- | --- | --- | --- | --- | --- | --- |
| 1 | M | 70 | 18 |  |  |  |  | 30 | 31 | 500 |
| 2 | M | 75 | 8 | 35 | 55 | 2 | 2.5 | 29 | 28 | 580 |
| 3 | M | 63 | 13 | 24 | 36 | 2 | 2 | 30 | 31 | 100 |
| 4 | M | 65 | 18 | 18 | 34 | 2 | 2 | 30 | 31 | 400 |
| 5 | M | 60 | 8 | 27 | 37.5 | 2 | 2 | 29 | 28 | 440 |
| 6 | M | 74 | 18 | 17 | 32 | 2 | 2.5 | 29 | 28 | 615 |
| 7 | F | 73 | 13 | 16 | 23 | 2 | 2 | 29 | 31 | 550 |
| 8 | M | 63 | 13 | 20 | 30 | 2 | 2 | 30 | 29 | 775 |
| 9 | M | 70 | 13 | 11 | 24 | 2 | 2 | 30 | 29 | 710 |
| 10 | M | 61 | 13 |  | 44 |  | 2.5 | 30 | 27 | 340 |
| 11 | M | 60 | 18 | 25 | 33 | 2 | 2 | 30 | 28 | 675 |
| 12 | M | 77 | 18 | 7 | 23 | 1.5 | 2 | 29 | 29 | 400 |
| 13 | M | 57 | 18 | 32 | 67 | 2 | 3 | 28 | 32 | 460 |
| 14 | M | 82 | 8 | 20 | 27 | 2 | 2 | 29 | 25 | 550 |
| 15 | M | 62 | 13 | 26 | 47 | 2 | 2.5 | 29 | 29 | 280 |
| **means** |  | 67 | 14 | 21 | 37 | 2 | 2 | 29 | 29 | 492 |

HCs were recruited for the study only if the following criteria were satisfied: (i) absence of neurological and/or psychiatric diseases, (ii) absence of psychological and/or cognitive disorders, (iii) absence of medications with psychotropic action and (iv) MMSE above or equal to 25. Patients were tested in three different experimental sessions over three different days. During the first one, they completed a neuropsychological assessment under their daily dopaminergic treatment. After this first session, patients were tested in two experimental conditions (in two different days), always at the same time of the day, with approximately 15 days as an intersession interval. In the OFF Condition, patients had been in drug withdrawal for 18 hours, and they were tested in the morning. In the ON Condition they were examined 60 min after their first morning therapeutic dose of levodopa and/or dopamine agonists. The Condition was counterbalanced across patients so that half of them performed the first experimental session in ON Condition and the other half in OFF. HCs performed the task in a single session, after which they were administered the MMSE. Moreover, to ascertain the efficacy of the dopaminergic medication in improving extrapyramidal motor symptoms, after each session, the patients were administered the UPDRS-Part III (Motor examination, Fahn and Elton, 1987) (a 27-items scale where each item is evaluated on a 5-point Likert scale, ranging from 0 to 4, where higher scores indicating higher extrapyramidal motor symptoms) and the Hoehn and Yahr scale (Hoehn and Yahr, 1967) (this scale identifies eight illness stages, indicated with the following numbers: 0-1-1.5-2-2.5-3-4-5, in which higher scores indicated higher extrapyramidal motor symptoms) scales in both ON and OFF conditions (see Table 1). The experimental protocol was approved by the ethics committee of the Fondazione Santa Lucia (Prot. CE/PROG.939) and was carried out in accordance with the ethical standards of the 1964 Declaration of Helsinki and later amendments. Participants gave their written informed consent to take part in the study. For indication about the Neuropsychological assessment performed see Supplementary Information.

### Experimental stimuli and set-up

The stimuli of the motor interaction task were the same used in previous studies (Sacheli et al., 2015a, b, 2018; Candidi et al., 2017; Gandolfo et al., 2019; Era et al., 2020a, 2020b; Moreau et al., 2020; Fini et al., 2021). They consisted of ten grasping movements performed by a virtual partner (five precision and five power grips). In detail, 3DS Max 2011 (Autodesk, Inc.) was employed to design the virtual scenario, where a virtual partner (created in Maya 2011 (Autodesk, Inc.) using a customized Python script (Prof. Orvalho V., Instituto de Telecomunicacoes, Porto University) was inserted. The virtual partner was created starting from the kinematics of a real person’s arm (recorded via a SMART-D motion capture system [Bioengineering Technology & Systems {B|T|S}]) that was recorded while grasping the upper part of a bottle (precision grip) or the lower part (power grip) with his right hand. Importantly, the virtual partner was displayed from the shoulders down, without the neck and head, to prevent facial expressions from exerting unwanted influences. Precision and power grip clips lasted the same amount of time (∼2000 ms). At the beginning of each stimulus, the virtual partner was still, with its hand positioned on the table. A variable amount of time (i.e., between 200 and 500 ms) after participants released the Start button (see below), the virtual partner started moving. The time of the virtual partner’s touch on the bottle was computed thanks to a photodiode, positioned on the screen where the virtual partner was displayed and detecting the appearance of a white square displayed on the frame where the virtual partner touched the bottle.

### Experimental task

Participants were requested to engage in a controlled motor interaction ‘Joint-Grasping task’ (Sacheli et al.,2015a, 2015b; Candidi et al., 2017; Gandolfo et al., 2019; Era et al., 2020a, 2020b; 2018; Boukarras et al., 2021; Moreau et al., 2020; Fini et al., 2021, Figure 1), where participants’ goal can only be achieved by predicting and monitoring the virtual partner’s movements and, therefore, adapting to them in real-time in order to grasp an object in synchrony. In detail, participants sat at a rectangular table, with a bottle-shaped object located in front of them. The object could be reached on two different parts: its lower part performing a whole-hand grasp (power grip) and its upper part performing a thumb-index precision grip (Movement Type factor). The virtual partner was presented on a monitor positioned on the table, behind the bottle-shaped object (Figure 1). Each trial started with the participant positioning their right hand over a start-button on the table, and their index finger and thumb touching each other. In different blocks, participants were required to: (i) monitor the movement of their partner online, in order to select whether to perform a precision grip or a power grasping based on the movement of the partner, as they were asked to perform an opposite or same action compared to that of their partner (Interactive condition) or (ii) follow an auditory instruction, indicating to perform a precision grip or power grasping regardless of what movement their partner performed (Cued condition), thus without the necessity to predict and monitor the partner’s actions. In both conditions, participants were instructed to grasp the bottle-shaped object as synchronously as possible with their virtual partner performing complementary or imitative actions with respect to it (Interaction type factor). In the imitative condition, participants grasped the same portion of the object as the virtual partner. In the complementary condition, instead, participants grasped the bottle-shaped object on different parts (e.g., the virtual partner grasped the lower part via power grip, and the participants grasped the upper part via precision grip, or vice-versa). In order to promote the need to monitor the virtual partner’s actions in the entire experiment, in 36% of trials the virtual partner performed a movement change by switching from a power grasp to a precision grip (or vice versa) during the reaching phase of the movement (Correction factor). In each session, participants performed two successive 64-trial Interactive and two successive 64-trial Cued blocks (in a counterbalanced order across participants) each comprising half power grasp and half precision grips. Thus, participants performed 10 NoCorrection and 6 Correction trials in each condition of the following 2×2×2 design: 2 (Interactive/Cued) × 2 (Complementary/Imitative) × 2 (Precision/Power grip). Stimuli presentation and randomization were controlled by E-Prime2 software (Psychology Software Tools Inc., Pittsburgh, PA). The experimental protocol never lasted more than 45 min to avoid exceeding the time-window of the effect of the dopaminergic therapy.

The moment in which participants touched the bottle was recorded thanks to touch-sensitive markers placed on the target spots of the object at 15 and 23 cm of the total height of the object. The trial timeline was as follows: participants heard the imitative/complementary or up/down auditory instruction, and they were presented with a still virtual partner. After its appearance, they were allowed to release the start button, and, upon releasing it, the virtual partner started its reach to -to-grasp movement. Thus, the beginning of the interaction and, particularly, of the movement of the virtual partner, was constrained to participants releasing the start button. This was done to overcome patients’ difficulty in initiating movements (present in patients with PD - Marsden, 1989) and allow them to engage in a real-time interaction with their partner. Movements were always performed with the right hand. The intertrial interval depended on the time participants took to go back from the bottle to the starting button. The experimenter manually moved to the following trial as soon as participants went back to the starting position and pressed the start button.

### Behavioural data

We excluded from the analyses the trials in which participants (i) missed the touch-sensitive markers, preventing from recording their responses, (ii) did not respect their Imitative/Complementary or Up/Down instructions, (iii) behavioural values that fell 2.5 SDs above or below each individual mean for each experimental condition (outlier trials) (on average, excluded trials of Grasping Asynchrony = 14.9 ± 10.2 – 5.47% of the total, specifically: 14 ± 10.4 for the PD ON group; 13.2 ± 8.5 for the PD OFF group; 17.6 ± 11.6 for the HCs group).

We considered as main behavioural measure Grasping Asynchrony (GAsynchr), i.e. the absolute value of time delay between the participant’s and virtual partner’s touch-time on the bottle-shaped object as the main behavioural measure. We subtracted from each individual’s mean in the Interactive conditions the corresponding individual’s means in the Cued conditions (as in Era et al., 2023). This way, we indexed the participants’ ability to perform the motor interaction task (requiring predicting and monitoring the virtual partner’s actions), net of their baseline ability in performing precision and power grasping actions as measured in the Cued condition (not requiring predicting and monitoring the virtual partner’s actions). To test whether the Condition factor (On/Off medication) influenced the Grasping Asynchrony in the Cued trials, before running the analysis on the subtraction (Interactive minus Cued), we run ran a t-test between Grasping Asynchrony in the ‘On’ and ‘Off’ Condition only in the Cued trials. Results showed no significant difference between the ‘On’ and ‘Off’ Conditions [t(14) = −1.66, pP = 0.12], indicating that the factor Condition had no effect on Grasping Asynchrony in the Cued trials.

We then subtracted from each individual’s mean of Grasping Asynchrony in the Interactive conditions the corresponding individual’s means in the Cued conditions. Detailed analyses of reaction times and movement times are provided in the Supplementary materials.

### EEG recording and preprocessing

EEG signals were recorded and amplified using a Neuroscan SynAmps RT amplifiers system (Compumedics Limited, Melbourne, Australia). These signals were acquired from 58 tin scalp electrodes embedded in a fabric cap (Electro-Cap International, Eaton, OH), arranged according to the 10-20 system. The EEG was recorded from the following channels: Fp1, Fpz, Fp2, AF3, AF4, F7, F5, F3, F1, Fz, F2, F4, F6, F8, FC5, FC3, FC1, FCz, FC2, FC4, FC6, T7, C5, C3, C1, Cz, C2, C4, C6, T8, TP7, CP5, CP3, CP1, CPz, CP2, CP4, CP6, TP8, P7, P5, P3, P1, Pz, P2, P4, P6, P8, PO7, PO3, AF7, POz, AF8, PO4, PO8, O1, Oz and O2. Horizontal electro-oculogram (HEOG) was recorded bipolarly from electrodes placed on the outer catchi of each eye and signals from the left earlobe were also recorded.

All electrodes were physically referenced to an electrode placed on the right earlobe and were algebraically re-referenced off-line to the average of both earlobe electrodes. Impedance was kept below 5 KΩ for all electrodes for the whole duration of the experiment, amplifier hardware band-pass filter was 0.1-200 Hz and sampling rate was 1000 Hz. Offline, we applied a high pass filter to 0.1 and a low pass filter to 48 in order to remove slow drifts and line noise (50 Hz) from the signal. The two filters were applied separately as suggested by the latest version of EEGLAB (Delorme et al., 2004). Then, channels exhibiting excessive amounts of noise were detected by running the clean_rawdata function as implemented in EEGLAB (Delorme et al., 2004), by removing the channels which trace correlated with their direct neighbours for less than 80% of the time. After removing the above mentioned channels, we epoched the continuous data into segments of 10.000 milliseconds – from −5.000 to 5.000 around the beginning of each trial. In order to identify and remove from the signal ocular artefacts (eye blinks and saccades) and excessive muscular noise, an Independent Component Analysis (ICA - Jung et al., 2000) was performed on the epoched data. Artefactual components were identified based on their time course and topography and removed. On average, 9 (± 2,74) components were removed per recording session. After the ICA, the bad electrodes previously removed were interpolated, with their signal being reconstructed based on the one from their surrounding neighbours with a “weighted” approach, as implemented in Fieldtrip (Oostenveld et al., 2011). Finally, EEG data were re-referenced to the average of all channels, and epochs containing remaining artefacts such as electrode jumps were removed following visual inspection.

### EEG Analysis

#### Univariate analysis

In order to investigate whether the neural correlates of action monitoring would be modulated in patients with PD, both comparing ON and OFF conditions (Condition factor), and comparing their performance to the one of by the control group (Group factor), we analysed the event-related potentials (ERPs) over fronto-central electrodes (i.e., FCz) time-locked to the correction of the VP (or the equivalent time frame in the No-Correction trials), as well as the activity in the time-frequency domain focused on midfrontal Delta-Theta.

#### ERPs

Before ERP averaging across subjects for each condition, EEG time-series were high-pass filtered at 1 Hz to remove slow drifts from the data, hence reducing the contribution of slow potentials (in previous works of our group with the same paradigm it was checked that grand average waveforms with different filters (i.e. 0.5, 1 Hz) maintain the same morphology, without introducing distorsions, see Moreau et al., (2020)). Since we found no ERN in response to the VP’s correction in any condition (see Discussion), only the Pe component was further analysed, being quantified as the mean amplitude in the time window between 200-450 ms over electrode FCz. No other components (including any trace of the late Pe over parietal sites - see Fig. S4) were found.

#### ERD/S

After preprocessing, as in the time-lock analysis, we high-pass filtered the segmented EEG signal at 1 Hz. Moreover, to remove the effects of ERPs in the time-frequency domain we subtracted the individual mean evoked response from each single trial, thus removing phase-locked activity (Sauseng et al., 2007; Pezzetta et. 2018). Each epoch was then put in the frequency domain using a Hanning-tapered window (Cohen, 2014) with a 50 ms time resolution, obtaining single-trial power representations for frequencies in a range from 1 to 40 Hz in steps of .25. The obtained induced power was averaged over trials for each condition and each subject, and the grand averages across subjects were displayed as event-related desynchronization/synchronization (ERD/ERS) with respect to a baseline period ranging from −.2 to 0 ms before the correction occurred.

To statistically compare midfrontal Delta-Theta across experimental factors, after visually inspecting the time-frequency plots we identified a coherent pattern of activity spanning from Delta (2-4 Hz) to Theta (4-7 Hz) (Fig. 4, 5, 6), thus we extracted the ERD/ERS for the Delta-Theta band (2-7 Hz) between 200 and 700 ms after the correction (or the corresponding frame in the NoCorrection trials) and analysed the modulation of power over FCz (as in Moreau et al., 2020).

Finally, we computed a difference in power, for each group, between Interactive and Cued trials. We run three separate ANOVAs comparing such differences in power between PD ON and OFF (with Group as within-subject factor), and between PD ON/OFF and HCs (with Group as between-subject factor). As a control, we also run all statistical analysis on the ERD/ERS for the Alpha (8-13 Hz) and Beta (14-30 Hz) band.

### Interaction between behavioural and EEG data

To test the influence of midfrontal Delta-Theta synchronization on participants’ ability to perform the Joint-Grasping task, we entered as continuous predictor single-trials data of midfrontal Delta-Theta ERS in a linear mixed model, performed with R Studio software. Single trials grasping Asynchrony scores were the dependent variable. We included Condition (Interactive, Cued) and Group (PD ON, OFF, HC) as categorical predictors. Statistical significance of fixed effects was determined using type III ANOVA test, with the lmer function from lme4 package. Estimates of slopes of the continuous predictor trend for each level of the factor were performed with the ‘Estimated Marginal Means’ R package (version 1.3.3, Lenth, 2017) via the emtrends functions.

The continuous predictor entered in the linear mixed model was mean-centered. The model included fixed effects for Condition and Group factors, the continuous predictor Delta-Theta ERS, and as random intercept the Participants and number of trials, as more complex models, including the random slope of the Condition factor did not converge.

### Multivariate analysis

To better identify the neural patterns that were sensitive to the interactive context (i.e., Interactive vs Cued), and as well as to investigate their modulation by PD and/or deficits in the levels of dopamine in the brain, we adopted a multivariate approach. Multivariate analyses are more sensitive than univariate analyses (Haxby et al., 2001; Tucciarelli et al., 2015), since they assume that the processing of different stimuli or tasks rely on different neural patterns to be exploited, thus they treat whole-brain sensor/source-level data as response patterns rather than investigating changes between average responses for each single sensor/source (as in the univariate approach) (Grootswagers et al., 2017a). Analyses were performed using the MVPA-Light toolbox (Treder, 2020) and custom scripts in Matlab R2019a. Prior to every classification, data were normalized using z-scores to center and scale the training data, providing numerical stability (Treder, 2020). Across all classification analyses, if a class had fewer trials than another, we corrected the imbalance by undersampling the over-represented condition (i.e., randomly removing trials). Then, using the preprocessing option mv_preprocess_average_samples, training data (i.e., EEG trials) from the same class (i.e., Interactive/Cued) were randomly split into 5 groups and averaged, so that the single-trial dataset was replaced by 5 averaged trials (i.e., samples). The classifications were then run using these averaged samples, as this step is known to increase the signal-to-noise ratio (Grootswagers et al., 2017; Treder, 2020).

We performed binary classification analyses in time (−4000 to 1000 ms around the VP’s correction), using Interactive/Cued samples, for each Group separately (i.e., HC, ON, and OFF). To do so, a Linear Discriminant Analysis (LDA) classifier was implemented using the MVPA-Light toolbox. For linearly separable data, an LDA classifier divides the data space into n regions, depending on the number of classes, and finds the optimal separating boundary between them using a discriminant function to find whether the data fall on the decision boundary (i.e., 50% classification accuracy) or far from it (i.e., > 50% classification accuracy). A k-fold cross-validation procedure in which samples were divided into 5 folds was repeated 5 times so that each sample was either used for training or testing at least once, and the raw output of the model was extracted and averaged for each decoded class at the subject-level. The decoding curves for each condition were then compared through 1) a dependent samples t-test across time between PD ON and OFF groups, and 2) two independent samples t-tests across time between PD ON/OFF and HCs groups. Statistical significance was assessed through non-parametric cluster-based permutations (Maris & Oostenveld, 2007).

Moreover, to investigate what electrodes contributed most to the classification over time, we performed a searchlight analysis using two neighboring matrices for time points and electrodes, respectively. Searchlight analysis is one approach to localize multivariate effects, as it strikes a balance between localization and statistical power (Kriegeskorte et al., 2006; Treder, 2020). Thus, in this analysis each electrode/time-point and its direct neighbors acted as features for the classification, resulting in a channels x time points matrix of accuracy scores tested for statistical significance. By plotting the results from this matrix also on a spatial topography averaged in specific time windows of interest, we then visualized which electrodes carried the most weight in the temporal decoding.

### Data handling and statistics

We run three different sets of ANOVAs on Grasping Asynchrony data (Interactive minus Cued Condition): (i) a within-participants ANOVA to compare the performance of patients with PD in the motor interactions task between ON and OFF conditions. This ANOVA had Condition (On/Off) × Interaction type (Complementary/Imitative) × Movement Type (Precision/Power grip) × Correction (Correction/NoCorrection) as within-subject factors; (ii) a mixed ANOVA, to compare the performance of PD ON patients and of HCs in the motor interactions task. This ANOVA had Group (PD ON/HC) as between-subjects factor and Interaction type (Complementary/Imitative) × Movement Type (Precision/Power grip) × Correction (Correction/NoCorrection) as within-subject factors; (iii) a mixed ANOVA to compare the performance of PD OFF and of HCs at the motor interactions task. This ANOVA had Group (PD OFF/HC) as between-subjects factor and Interaction type (Complementary/Imitative) × Movement Type (Precision/Power grip) × Correction (Correction/NoCorrection) as within-subject factors. The same ANOVAs (including the Interactivity factor) were performed to compare ERPs mean amplitude values and ERD/ERS values across conditions. Since our main hypothesis did not focus on Interaction Type (Complementary/Imitative) and Movement Type (Precision/Power), these factors were collapsed in order to have a higher number of trials for each condition. All tests of significance were based on an level of 0.05. Post hoc tests were performed using the Newman–Keuls method when appropriate. All statistical analyses were performed in Matlab R2019a and using the JASP Software (JASP Team (2023)). In all the analyses, we report significant main effects or interactions that include the factor Condition and Group. All other main effects and interactions are reported in Supplementary Tables 1-16.

## REFERENCES

Blesa, J., Foffani, G., Dehay, B., Bezard, E., & Obeso, J. A. (2021). Motor and non-motor circuit disturbances in early Parkinson disease: which happens first? Nature Reviews. Neuroscience, 23(2), 115–128. 10.1038/s41583-021-00542-9

Boukarras, S., Era, V., Aglioti, S. M., & Candidi, M. (2021). Competence-based social status and implicit preference modulate the ability to coordinate during a joint grasping task. Scientific Reports, 11(1). 10.1038/s41598-021-84280-z

Boukarras, S., Özkan, D. G., Era, V., Moreau, Q., Tieri, G., & Candidi, M. (2022). Midfrontal theta transcranial alternating current stimulation facilitates motor coordination in dyadic Human–Avatar interactions. Journal of Cognitive Neuroscience, 34(5), 897–915. 10.1162/jocn_a_01834

Campbell, J. I. D., & Thompson, V. A. (2012). MorePower 6.0 for ANOVA with relational confidence intervals and Bayesian analysis. Behavior Research Methods, 44(4), 1255–1265. 10.3758/s13428-012-0186-0

Candidi, M., Sacheli, L. M., Era, V., Canzano, L., Tieri, G., & Aglioti, S. M. (2017). Come together: human–avatar on-line interactions boost joint-action performance in apraxic patients. Social Cognitive and Affective Neuroscience, 12(11), 1793–1802. 10.1093/scan/nsx114

Cavanagh, J. F., & Frank, M. J. (2014). Frontal theta as a mechanism for cognitive control. Trends in Cognitive Sciences, 18(8), 414–421. 10.1016/j.tics.2014.04.012

Cohen, M. X. (2011). Error-related medial frontal theta activity predicts cingulate-related structural connectivity. NeuroImage, 55(3), 1373–1383. 10.1016/j.neuroimage.2010.12.072

Cohen, M. X. (2014). Analyzing neural Time series data. In The MIT Press eBooks. 10.7551/mitpress/9609.001.0001

De Bondt, C. C., Gerrits, N. J. H. M., Veltman, D. J., Berendse, H. W., Van Den Heuvel, O. A., & Van Der Werf, Y. D. (2016). Reduced task-related functional connectivity during a set-shifting task in unmedicated early-stage Parkinson’s disease patients. BMC Neuroscience, 17(1). 10.1186/s12868-016-0254-y

Delorme, A., & Makeig, S. (2004). EEGLAB: an open source toolbox for analysis of single-trial EEG dynamics including independent component analysis. Journal of Neuroscience Methods, 134(1), 9–21. 10.1016/j.jneumeth.2003.10.009

Endrass, T., & Ullsperger, M. (2014). Specificity of performance monitoring changes in obsessive-compulsive disorder. Neuroscience & Biobehavioral Reviews/Neuroscience and Biobehavioral Reviews, 46, 124–138. 10.1016/j.neubiorev.2014.03.024

Era, V., Aglioti, S. M., & Candidi, M. (2019). Inhibitory Theta Burst Stimulation Highlights the Role of Left aIPS and Right TPJ during Complementary and Imitative Human–Avatar Interactions in Cooperative and Competitive Scenarios. Cerebral Cortex, 30(3), 1677–1687. 10.1093/cercor/bhz195

Era, V., Aglioti, S. M., Mancusi, C., & Candidi, M. (2018). Visuo-motor interference with a virtual partner is equally present in cooperative and competitive interactions. Psychological Research, 84(3), 810–822. 10.1007/s00426-018-1090-8

Era, V., Boukarras, S., & Candidi, M. (2019). Neural correlates of action monitoring and mutual adaptation during interpersonal motor coordination. Physics of Life Reviews, 28, 43–45. 10.1016/j.plrev.2019.01.022

Era, V., Candidi, M., Gandolfo, M., Sacheli, L. M., & Aglioti, S. M. (2018). Inhibition of left anterior intraparietal sulcus shows that mutual adjustment marks dyadic joint-actions in humans. Social Cognitive and Affective Neuroscience, 13(5), 492–500. 10.1093/scan/nsy022

Era, V., Candidi, M., Pezzetta, R., Pulcini, C., D’Antonio, S., Zabberoni, S., Peppe, A., Costa, A., Taglieri, S., Carlesimo, G. A., & Aglioti, S. M. (2022). The dopaminergic system supports flexible and rewarding dyadic motor interactive behaviour in Parkinson’s Disease. Social Cognitive and Affective Neuroscience, 18(1). 10.1093/scan/nsac040

Falkenstein, M., Hielscher, H., Dziobek, I., Schwarzenau, P., Hoormann, J., Sundermann, B., & Hohnsbein, J. (2001). Action monitoring, error detection, and the basal ganglia: an ERP study. NeuroReport/Neuroreport, 12(1), 157–161. 10.1097/00001756-200101220-00039

Falkenstein, M., Hohnsbein, J., Hoormann, J., & Blanke, L. (1991). Effects of crossmodal divided attention on late ERP components. II. Error processing in choice reaction tasks. Electroencephalography and Clinical Neurophysiology, 78(6), 447–455. 10.1016/0013-4694(91)90062-9

Fini, C., Era, V., Da Rold, F., Candidi, M., & Borghi, A. M. (2021). Abstract concepts in interaction: the need of others when guessing abstract concepts smooths dyadic motor interactions. Royal Society Open Science, 8(7), 201205. 10.1098/rsos.201205

Fish, J. (2011). Unified Parkinson’s disease rating scale. In Springer eBooks (pp. 2576–2577). 10.1007/978-0-387-79948-3_1836

Fusco, G., Cristiano, A., Perazzini, A., & Aglioti, S. M. (2022). Neuromodulating the performance monitoring network during conflict and error processing in healthy populations: Insights from transcranial electric stimulation studies. Frontiers in Integrative Neuroscience, 16. 10.3389/fnint.2022.953928

Fusco, G., Fusaro, M., & Aglioti, S. M. (2020). Midfrontal-occipital θ-tACS modulates cognitive conflicts related to bodily stimuli. Social Cognitive and Affective Neuroscience, 17(1), 91–100. 10.1093/scan/nsaa125

Galea, J. M., Bestmann, S., Beigi, M., Jahanshahi, M., & Rothwell, J. C. (2012). Action reprogramming in Parkinson’s disease: response to prediction error is modulated by levels of dopamine. the Journal of Neuroscience/the Journal of Neuroscience, 32(2), 542–550. 10.1523/jneurosci.3621-11.2012

Gandolfo, M., Era, V., Tieri, G., Sacheli, L. M., & Candidi, M. (2019). Interactor’s body shape does not affect visuo-motor interference effects during motor coordination. Acta Psychologica, 196, 42–50. 10.1016/j.actpsy.2019.04.003

Grootswagers, T., Wardle, S. G., & Carlson, T. A. (2017). Decoding Dynamic Brain Patterns from Evoked Responses: A Tutorial on Multivariate Pattern Analysis Applied to Time Series Neuroimaging Data. Journal of Cognitive Neuroscience, 29(4), 677–697. 10.1162/jocn_a_01068

Haxby, J. V., Gobbini, M. I., Furey, M. L., Ishai, A., Schouten, J. L., & Pietrini, P. (2001). Distributed and overlapping representations of faces and objects in ventral temporal cortex. Science, 293(5539), 2425–2430. 10.1126/science.1063736

Hoehn, M. M., & Yahr, M. D. (1967). Parkinsonism. Neurology, 17(5), 427. 10.1212/wnl.17.5.427

Holroyd, C. B., & Coles, M. G. H. (2002). The neural basis of human error processing: Reinforcement learning, dopamine, and the error-related negativity. Psychological Review, 109(4), 679–709. 10.1037/0033-295x.109.4.679

Jocham, G., & Ullsperger, M. (2009). Neuropharmacology of performance monitoring. Neuroscience & Biobehavioral Reviews/Neuroscience and Biobehavioral Reviews, 33(1), 48–60. 10.1016/j.neubiorev.2008.08.011

Jung, T., Makeig, S., Westerfield, M., Townsend, J., Courchesne, E., & Sejnowski, T. J. (2000). Removal of eye activity artifacts from visual event-related potentials in normal and clinical subjects. Clinical Neurophysiology, 111(10), 1745–1758. 10.1016/s1388-2457(00)00386-2

Kerns, J. G., Cohen, J. D., MacDonald, A. W., Cho, R. Y., Stenger, V. A., & Carter, C. S. (2004). Anterior cingulate conflict monitoring and adjustments in control. Science, 303(5660), 1023–1026. 10.1126/science.1089910

Klein, T. A., Ullsperger, M., & Danielmeier, C. (2013). Error awareness and the insula: links to neurological and psychiatric diseases. Frontiers in Human Neuroscience, 7. 10.3389/fnhum.2013.00014

Kricheldorff, J., Ficke, J., Debener, S., & Witt, K. (2023). Impaired proactive cognitive control in Parkinson’s disease. Brain Communications, 5(6). 10.1093/braincomms/fcad327

Liu, Y., Gehring, W. J., Weissman, D. H., Taylor, S. F., & Fitzgerald, K. D. (2012). Trial-by-Trial Adjustments of Cognitive Control Following Errors and Response Conflict are Altered in Pediatric Obsessive Compulsive Disorder. Frontiers in Psychiatry, 3. 10.3389/fpsyt.2012.00041

Luu, P., Tucker, D. M., & Makeig, S. (2004). Frontal midline theta and the error-related negativity: neurophysiological mechanisms of action regulation. Clinical Neurophysiology, 115(8), 1821–1835. 10.1016/j.clinph.2004.03.031

Maris, E., & Oostenveld, R. (2007). Nonparametric statistical testing of EEG- and MEG-data. Journal of Neuroscience Methods, 164(1), 177–190. 10.1016/j.jneumeth.2007.03.024

Moreau, Q., Candidi, M., Era, V., Tieri, G., & Aglioti, S. M. (2020). Midline frontal and occipito-temporal activity during error monitoring in dyadic motor interactions. Cortex, 127, 131–149. 10.1016/j.cortex.2020.01.020

Moreau, Q., Tieri, G., Era, V., Aglioti, S. M., & Candidi, M. (2022). The performance monitoring system is attuned to others’ actions during dyadic motor interactions. Cerebral Cortex, 33(1), 222–234. 10.1093/cercor/bhac063

Oehrn, C. R., Hanslmayr, S., Fell, J., Deuker, L., Kremers, N. A., Lam, A. T. D., Elger, C. E., & Axmacher, N. (2014). Neural communication patterns underlying conflict detection, resolution, and adaptation. the Journal of Neuroscience/the Journal of Neuroscience, 34(31), 10438–10452. 10.1523/jneurosci.3099-13.2014

Oostenveld, R., Fries, P., Maris, E., & Schoffelen, J. (2011). FieldTrip: open source software for advanced analysis of MEG, EEG, and invasive electrophysiological data. Computational Intelligence and Neuroscience, 2011, 1–9. 10.1155/2011/156869

Overbeek, T. J., Nieuwenhuis, S., & Ridderinkhof, K. R. (2005). Dissociable components of error processing. Journal of Psychophysiology, 19(4), 319–329. 10.1027/0269-8803.19.4.319

Parker, K. L., Chen, K., Kingyon, J. R., Cavanagh, J. F., & Narayanan, N. S. (2015). Medial frontal 14-Hz activity in humans and rodents is attenuated in PD patients and in rodents with cortical dopamine depletion. Journal of Neurophysiology, 114(2), 1310–1320. 10.1152/jn.00412.2015

Pezzetta, R., Ozkan, D. G., Era, V., Tieri, G., Zabberoni, S., Taglieri, S., Costa, A., Peppe, A., Caltagirone, C., & Aglioti, S. M. (2023). Combined EEG and immersive virtual reality unveil dopaminergic modulation of error monitoring in Parkinson’s Disease. NPJ Parkinson’s Disease, 9(1). 10.1038/s41531-022-00441-5

Ponsi, G., Scattolin, M., Villa, R., & Aglioti, S. M. (2021). Human moral decision-making through the lens of Parkinson’s disease. NPJ Parkinson’s Disease, 7(1). 10.1038/s41531-021-00167-w

Ridderinkhof, K. R., Ramautar, J. R., & Wijnen, J. G. (2009). To PE or not to PE: A P3-like ERP component reflecting the processing of response errors. Psychophysiology, 46(3), 531–538. 10.1111/j.1469-8986.2009.00790.x

Sacheli, L. M., Candidi, M., Era, V., & Aglioti, S. M. (2015). Causative role of left aIPS in coding shared goals during human–avatar complementary joint actions. Nature Communications, 6(1). 10.1038/ncomms8544

Sacheli, L. M., Christensen, A., Giese, M. A., Taubert, N., Pavone, E. F., Aglioti, S. M., & Candidi, M. (2015). Prejudiced interactions: implicit racial bias reduces predictive simulation during joint action with an out-group avatar. Scientific Reports, 5(1). 10.1038/srep08507

Sacheli, L. M., Diana, L., Ravani, A., Beretta, S., Bolognini, N., & Paulesu, E. (2023). Neuromodulation of the Le t Inferior Frontal Cortex Affects Social Monitoring during Motor Interactions. Journal of Cognitive Neuroscience, 35(11), 1788–1805. 10.1162/jocn_a_02046

Sacheli, L. M., Musco, M. A., Zazzera, E., & Paulesu, E. (2021). Mechanisms for mutual support in motor interactions. Scientific Reports, 11(1). 10.1038/s41598-021-82138-y

Sacheli, L. M., Tieri, G., Aglioti, S. M., & Candidi, M. (2018). Transitory inhibition of the left anterior intraparietal sulcus impairs joint actions: a continuous Theta-Burst stimulation study. Journal of Cognitive Neuroscience, 30(5), 737–751. 10.1162/jocn_a_01227

Sauseng, P., Klimesch, W., Gruber, W., Hanslmayr, S., Freunberger, R., & Doppelmayr, M. (2007). Are event-related potential components generated by phase resetting of brain oscillations? A critical discussion. Neuroscience, 146(4), 1435–1444. 10.1016/j.neuroscience.2007.03.014

Sebanz, N., Bekkering, H., & Knoblich, G. (2006). Joint action: bodies and minds moving together. Trends in Cognitive Sciences, 10(2), 70–76. 10.1016/j.tics.2005.12.009

Solié, C., Girard, B., Righetti, B., Tapparel, M., & Bellone, C. (2021). VTA dopamine neuron activity encodes social interaction and promotes reinforcement learning through social prediction error. Nature Neuroscience, 25(1), 86–97. 10.1038/s41593-021-00972-9

Spinelli, G., Pezzetta, R., Canzano, L., Tidoni, E., & Aglioti, S. M. (2022). Brain dynamics of Action Monitoring in Higher-Order Motor Control Disorders: the case of Apraxia. ENeuro, 9(2), ENEURO.0334-20.2021. 10.1523/eneuro.0334-20.2021

Treder, M. S. (2020). MVPA-Light: A Classification and Regression Toolbox for Multi-Dimensional Data. Frontiers in Neuroscience, 14. 10.3389/fnins.2020.00289

Tucciarelli, R., Turella, L., Oosterhof, N. N., Weisz, N., & Lingnau, A. (2015). MEG multivariate analysis reveals early abstract action representations in the lateral occipitotemporal cortex. the Journal of Neuroscience/the Journal of Neuroscience, 35(49), 16034–16045. 10.1523/jneurosci.1422-15.2015

Ullsperger, M., Danielmeier, C., & Jocham, G. (2014). Neurophysiology of Performance monitoring and Adaptive Behavior. Physiological Reviews, 94(1), 35–79. 10.1152/physrev.00041.2012

Ullsperger, M., & Von Cramon, D. Y. (2006). The role of intact frontostriatal circuits in error processing. Journal of Cognitive Neuroscience, 18(4), 651–664. 10.1162/jocn.2006.18.4.651

Wylie, S. A., Ridderinkhof, K. R., Elias, W. J., Frysinger, R. C., Bashore, T. R., Downs, K. E., Van Wouwe, N. C., & Van Den Wildenberg, W. P. M. (2010). Subthalamic nucleus stimulation influences expression and suppression of impulsive behaviour in Parkinson’s disease. Brain, 133(12), 3611–3624. 10.1093/brain/awq239

