## Supplementary Materials for "Dopaminergic modulation of behavioral and electrocortical markers of interpersonal performance monitoring in Parkinson’s Disease: insights from multivariate and univariate analyses"

**Neuropsychological and motor assessment**

Patients with PD received a complete neuropsychological assessment during the “On” Condition. In Supplemetary Table 17 we report the results from the sub-set of tests that were completed by all patients, including at least one test for each cognitive domain: General cognitive functioning: Mini Mental State Examination (MMSE; Measso et al., 1991) and Mini Mental Parkinson State Examination (MMPSE; Costa et al., 2013); Attention and Short Term Memory: Digit Span and Corsi Block tapping test forward (Monaco et al, 2013), Trail Making Test (subtest A, Giovagnoli et al, 1996); Executive functions: Stroop test (Barbarotto et al., 1998), Trail Making Test (subtest B, Giovagnoli et al, 1996), Modified Card Sorting Test (MCST, Nocentini et al, 2002), Digit Span and Corsi block tapping test backward (Monaco et al, 2013); Phonological, Semantic and Alternate Fluency (Costa et al 2014); Episodic Memory: immediate and delayed recall of a 15 words-list (Carlesimo et al., 1996), immediate and delayed prose recall (Carlesimo et al, 2002); Language: Objects and Verbs naming (BADA, Capasso e Miceli, 2001); Logic reasoning: Coloured Raven’s progressive matrices (Carlesimo et al, 1996); Visuo-spatial abilities: Copy of Rey’s Figure (Carlesimo et al., 2002), copy of drawing with and without landmarks (Carlesimo et al, 1996).

**Behavioural results – Movement Time**

In the 2 Condition (PD ON/OFF) × Interaction type (Complementary/Imitative) × Movement Type (Precision/Power grip) × Correction (Correction/NoCorrection), within-participants ANOVA on Movement Time only the main effect of the Interaction Type reached statistical significance [F(1, 14) = 20.006 , p = <.001, ηp² = .588] showing that patients’ movement time increased when performing an Complementary (M = 459.29 ms, SD = 203.38) rather than an Imitative (M = 398.21 ms, SD = 216.5) movement. As the primary goal of the present study was to assess any modulation of the behavioural performance by the Condition factor, we report all other interactions between factors in Supplementary Table 4.

In the 2 Group (PD ON/HC) x 2 Interaction type (Complementary/Imitative) × 2 Movement Type (Precision/Power grip) × 2 Correction (Correction/NoCorrection) mixed ANOVA on Movement Time the Correction and Interaction Type within-subjects factors reached statistical significance. In detail, the Interaction type factor [F(1, 14) = 14.174 , p < .001, ηp² = .336], showed that participants’ movement time increased when performing a Complementary (M = 421.98 ms, SD = 256.48) rather than an Imitative (M = 372.12 ms, SD = 253.41) movement. The Correction factor [F(1, 14) = 4.223, p = .049, ηp² = .131] highlighted that participants’ movement took longer when a Correction of the VP occurred (M = 413.81 ms, SD = 242.69) compared to when it did not (M = 382.29 ms, SD = 267.87). The Group factor did not interact with any within-subject factor, whereas all other interactions between factors are reported in Supplementary Table 5.

In the 2 Group (PD OFF/HC) x 2 Interaction type (Complementary/Imitative) × 2 Movement Type (Precision/Power grip) × 2 Correction (Correction/NoCorrection) mixed ANOVA on Movement Time the Correction and Interaction Type within-subjects factors reached statistical significance. In detail, the Interaction type factor [F(1, 14) = 10.266 , p = .003, ηp² = .268], showed that participants’ movement time increased when performing a Complementary (M = 429.20 ms, SD = 235.17) rather than an Imitative (M = 389.97 ms, SD = 218.64) movement. The Correction factor [F(1, 14) = 6.069, p = .020, ηp² = .178] highlighted that participants’ movement took longer when a Correction of the VP occurred (M = 426.98 ms, SD = 214.31) compared to when it did not (M = 392.20 ms, SD = 239.48). The Group factor did not interact with any within-subject factor, whereas all other interactions between factors are reported in Supplementary Table 6.

**Behavioural results – Reaction Times**

In the 2 Condition (PD ON/OFF) × Interaction type (Complementary/Imitative) × Movement Type (Precision/Power grip) × Correction (Correction/NoCorrection), within-participants ANOVA on Reaction Times no factor nor interaction between factors reached statistical significance.

In the 2 Group (PD ON/HC) x 2 Interaction type (Complementary/Imitative) × 2 Movement Type (Precision/Power grip) × 2 Correction (Correction/NoCorrection) mixed ANOVA on Reaction Times the Grip within-subjects factor reached statistical significance [F(1, 14) = 8.898 , p = .006, ηp² = .241]. In detail, participants’ reaction time in starting the movement increased when performing a Power (M = 91.66 ms, SD = 271.44) rather than a Precision(M = 53.04 ms, SD = 271.62) movement. The Group factor did not interact with any within-subject factor, and no other interactions between factors occurred.

In the 2 Group (PD OFF/HC) x 2 Interaction type (Complementary/Imitative) × 2 Movement Type (Precision/Power grip) × 2 Correction (Correction/NoCorrection) mixed ANOVA on Reaction Times no factor nor interaction between factors reached statistical significance.

All statistical results on Reaction Times are reported in Supplementary Table 7, 8 and 9.

| **SUPPLEMENTARY TABLE 1a – Grasping Asynchrony - Within-participants ANOVA PD ON vs PD OFF** | | | | | | | | | | | | |
| --- | --- | --- | --- | --- | --- | --- | --- | --- | --- | --- | --- | --- |
| **Within Subjects Effects** | | | | | | | | | | | | |
| **Cases** | | **Sum of Squares** | | **df** | | **Mean Square** | | **F** | | **p** | | **η²_p_** |
| Condition |  | 325514.420 |  | 1 |  | 325514.420 |  | 5.876 |  | 0.029 |  | 0.296 |
| Residuals |  | 775531.244 |  | 14 |  | 55395.089 |  |  |  |  |  |  |
| Interaction type |  | 123857.498 |  | 1 |  | 123857.498 |  | 12.707 |  | 0.003 |  | 0.476 |
| Residuals |  | 136456.067 |  | 14 |  | 9746.862 |  |  |  |  |  |  |
| Movement type |  | 148304.204 |  | 1 |  | 148304.204 |  | 11.762 |  | 0.004 |  | 0.457 |
| Residuals |  | 176516.405 |  | 14 |  | 12608.315 |  |  |  |  |  |  |
| Correction |  | 160748.022 |  | 1 |  | 160748.022 |  | 9.286 |  | 0.009 |  | 0.399 |
| Residuals |  | 242344.779 |  | 14 |  | 17310.341 |  |  |  |  |  |  |
| Condition ✻ Interaction type |  | 275.959 |  | 1 |  | 275.959 |  | 0.051 |  | 0.825 |  | 0.004 |
| Residuals |  | 75854.343 |  | 14 |  | 5418.167 |  |  |  |  |  |  |
| Condition ✻ Movement type |  | 1256.270 |  | 1 |  | 1256.270 |  | 0.114 |  | 0.741 |  | 0.008 |
| Residuals |  | 154791.623 |  | 14 |  | 11056.545 |  |  |  |  |  |  |
| Interaction type ✻ Movement type |  | 26385.249 |  | 1 |  | 26385.249 |  | 2.890 |  | 0.111 |  | 0.171 |
| Residuals |  | 127825.752 |  | 14 |  | 9130.411 |  |  |  |  |  |  |
| Condition ✻ Correction |  | 6037.006 |  | 1 |  | 6037.006 |  | 2.310 |  | 0.151 |  | 0.142 |
| Residuals |  | 36593.629 |  | 14 |  | 2613.831 |  |  |  |  |  |  |
| Interaction type ✻ Correction |  | 8987.412 |  | 1 |  | 8987.412 |  | 1.020 |  | 0.330 |  | 0.068 |
| Residuals |  | 123314.641 |  | 14 |  | 8808.189 |  |  |  |  |  |  |
| Movement type ✻ Correction |  | 2665.071 |  | 1 |  | 2665.071 |  | 0.923 |  | 0.353 |  | 0.062 |
| Residuals |  | 40414.687 |  | 14 |  | 2886.763 |  |  |  |  |  |  |
| Condition ✻ Interaction type ✻ Movement type |  | 722.310 |  | 1 |  | 722.310 |  | 0.180 |  | 0.678 |  | 0.013 |
| Residuals |  | 56115.215 |  | 14 |  | 4008.230 |  |  |  |  |  |  |
| Condition ✻ Interaction type ✻ Correction |  | 4.443 |  | 1 |  | 4.443 |  | 0.001 |  | 0.973 |  | 8.720×10^-5^ |
| Residuals |  | 50946.501 |  | 14 |  | 3639.036 |  |  |  |  |  |  |
| Condition ✻ Movement type ✻ Correction |  | 1708.523 |  | 1 |  | 1708.523 |  | 0.368 |  | 0.554 |  | 0.026 |
| Residuals |  | 65010.853 |  | 14 |  | 4643.632 |  |  |  |  |  |  |
| Interaction type ✻ Movement type ✻ Correction |  | 260502.621 |  | 1 |  | 260502.621 |  | 10.467 |  | 0.006 |  | 0.428 |
| Residuals |  | 348436.263 |  | 14 |  | 24888.304 |  |  |  |  |  |  |
| Condition ✻ Interaction type ✻ Movement type ✻ Correction |  | 5157.614 |  | 1 |  | 5157.614 |  | 0.607 |  | 0.449 |  | 0.042 |
| Residuals |  | 119020.391 |  | 14 |  | 8501.456 |  |  |  |  |  |  |
| *Note.*  Type III Sum of Squares | | | | | | | | | | | | |

| **SUPPLEMENTARY TABLE 1b – Grasping Asynchrony - Post Hoc Comparisons - Interaction type ✻ Movement type ✻ Correction** | | | | | | | | | | |
| --- | --- | --- | --- | --- | --- | --- | --- | --- | --- | --- |
|  | |  | | **Mean Difference** | | **SE** | | **t** | | **p_bonf_** |
| Complementary, Precision, Corr |  | Imitative, Precision, Corr |  | 102.595 |  | 29.601 |  | 3.466 |  | 0.033 |
|  |  | Complementary, Power, Corr |  | 1.869 |  | 28.727 |  | 0.065 |  | 1.000 |
|  |  | Imitative, Power, Corr |  | 14.621 |  | 23.822 |  | 0.614 |  | 1.000 |
|  |  | Complementary, Precision, NoCorr |  | 136.556 |  | 29.970 |  | 4.556 |  | 0.001 |
|  |  | Imitative, Precision, NoCorr |  | 82.889 |  | 25.519 |  | 3.248 |  | 0.062 |
|  |  | Complementary, Power, NoCorr |  | -6.688 |  | 28.242 |  | -0.237 |  | 1.000 |
|  |  | Imitative, Power, NoCorr |  | 113.370 |  | 32.801 |  | 3.456 |  | 0.032 |
| Imitative, Precision, Corr |  | Complementary, Power, Corr |  | -100.725 |  | 23.822 |  | -4.228 |  | 0.003 |
|  |  | Imitative, Power, Corr |  | -87.973 |  | 28.727 |  | -3.062 |  | 0.110 |
|  |  | Complementary, Precision, NoCorr |  | 33.961 |  | 25.519 |  | 1.331 |  | 1.000 |
|  |  | Imitative, Precision, NoCorr |  | -19.706 |  | 29.970 |  | -0.657 |  | 1.000 |
|  |  | Complementary, Power, NoCorr |  | -109.282 |  | 32.801 |  | -3.332 |  | 0.046 |
|  |  | Imitative, Power, NoCorr |  | 10.775 |  | 28.242 |  | 0.382 |  | 1.000 |
| Complementary, Power, Corr |  | Imitative, Power, Corr |  | 12.752 |  | 29.601 |  | 0.431 |  | 1.000 |
|  |  | Complementary, Precision, NoCorr |  | 134.686 |  | 28.242 |  | 4.769 |  | < .001 |
|  |  | Imitative, Precision, NoCorr |  | 81.020 |  | 32.801 |  | 2.470 |  | 0.476 |
|  |  | Complementary, Power, NoCorr |  | -8.557 |  | 29.970 |  | -0.286 |  | 1.000 |
|  |  | Imitative, Power, NoCorr |  | 111.500 |  | 25.519 |  | 4.369 |  | 0.002 |
| Imitative, Power, Corr |  | Complementary, Precision, NoCorr |  | 121.934 |  | 32.801 |  | 3.717 |  | 0.014 |
|  |  | Imitative, Precision, NoCorr |  | 68.268 |  | 28.242 |  | 2.417 |  | 0.538 |
|  |  | Complementary, Power, NoCorr |  | -21.309 |  | 25.519 |  | -0.835 |  | 1.000 |
|  |  | Imitative, Power, NoCorr |  | 98.748 |  | 29.970 |  | 3.295 |  | 0.057 |
| Complementary, Precision, NoCorr |  | Imitative, Precision, NoCorr |  | -53.666 |  | 29.601 |  | -1.813 |  | 1.000 |
|  |  | Complementary, Power, NoCorr |  | -143.243 |  | 28.727 |  | -4.986 |  | < .001 |
|  |  | Imitative, Power, NoCorr |  | -23.186 |  | 23.822 |  | -0.973 |  | 1.000 |
| Imitative, Precision, NoCorr |  | Complementary, Power, NoCorr |  | -89.577 |  | 23.822 |  | -3.760 |  | 0.013 |
|  |  | Imitative, Power, NoCorr |  | 30.481 |  | 28.727 |  | 1.061 |  | 1.000 |
| Complementary, Power, NoCorr |  | Imitative, Power, NoCorr |  | 120.058 |  | 29.601 |  | 4.056 |  | 0.006 |
| *Note.*  P-value adjusted for comparing a family of 28 | | | | | | | | | | |
| *Note.*  Results are averaged over the levels of: Condition | | | | | | | | | | |

| **SUPPLEMENTARY TABLE 2a – Grasping Asynchrony - Mixed ANOVA PD ON vs HC** | | | | | | | | | | | | | | | | | | | | | | | |
| --- | --- | --- | --- | --- | --- | --- | --- | --- | --- | --- | --- | --- | --- | --- | --- | --- | --- | --- | --- | --- | --- | --- | --- |
| **Within Subjects Effects** | | | | | | | | | | | | | | | | | | | | | | | |
| **Cases** | | | | | | | | | **Sum of Squares** | | | | | | **df** | | **Mean Square** | | **F** | | **p** | | **η²_p_** |
| Interaction type | | | | | | | |  | 88548.735 | | | | |  | 1 |  | 88548.735 |  | 21.479 |  | < .001 |  | 0.434 |
| Interaction type ✻ GROUP | | | | | | | |  | 5037.390 | | | | |  | 1 |  | 5037.390 |  | 1.222 |  | 0.278 |  | 0.042 |
| Residuals | | | | | | | |  | 115429.624 | | | | |  | 28 |  | 4122.487 |  |  |  |  |  |  |
| Movement type | | | | | | | |  | 165986.703 | | | | |  | 1 |  | 165986.703 |  | 9.402 |  | 0.005 |  | 0.251 |
| Movement type ✻ GROUP | | | | | | | |  | 3335.722 | | | | |  | 1 |  | 3335.722 |  | 0.189 |  | 0.667 |  | 0.007 |
| Residuals | | | | | | | |  | 494322.648 | | | | |  | 28 |  | 17654.380 |  |  |  |  |  |  |
| Correction | | | | | | | |  | 170940.743 | | | | |  | 1 |  | 170940.743 |  | 17.021 |  | < .001 |  | 0.378 |
| Correction ✻ GROUP | | | | | | | |  | 8138.574 | | | | |  | 1 |  | 8138.574 |  | 0.810 |  | 0.376 |  | 0.028 |
| Residuals | | | | | | | |  | 281201.414 | | | | |  | 28 |  | 10042.908 |  |  |  |  |  |  |
| Interaction type ✻ Movement type | | | | | | | |  | 21784.536 | | | | |  | 1 |  | 21784.536 |  | 3.533 |  | 0.071 |  | 0.112 |
| Interaction type ✻ Movement type ✻ GROUP | | | | | | | |  | 1740.170 | | | | |  | 1 |  | 1740.170 |  | 0.282 |  | 0.599 |  | 0.010 |
| Residuals | | | | | | | |  | 172654.823 | | | | |  | 28 |  | 6166.244 |  |  |  |  |  |  |
| Interaction type ✻ Correction | | | | | | | |  | 0.678 | | | | |  | 1 |  | 0.678 |  | 1.035×10^-4^ |  | 0.992 |  | 3.697×10^-6^ |
| Interaction type ✻ Correction ✻ GROUP | | | | | | | |  | 9551.789 | | | | |  | 1 |  | 9551.789 |  | 1.458 |  | 0.237 |  | 0.049 |
| Residuals | | | | | | | |  | 183447.625 | | | | |  | 28 |  | 6551.701 |  |  |  |  |  |  |
| Movement type ✻ Correction | | | | | | | |  | 1343.898 | | | | |  | 1 |  | 1343.898 |  | 0.199 |  | 0.659 |  | 0.007 |
| Movement type ✻ Correction ✻ GROUP | | | | | | | |  | 16800.783 | | | | |  | 1 |  | 16800.783 |  | 2.490 |  | 0.126 |  | 0.082 |
| Residuals | | | | | | | |  | 188938.898 | | | | |  | 28 |  | 6747.818 |  |  |  |  |  |  |
| Interaction type ✻ Movement type ✻ Correction | | | | | | | |  | 519471.682 | | | | |  | 1 |  | 519471.682 |  | 39.209 |  | < .001 |  | 0.583 |
| Interaction type ✻ Movement type ✻ Correction ✻ GROUP | | | | | | | |  | 19191.318 | | | | |  | 1 |  | 19191.318 |  | 1.449 |  | 0.239 |  | 0.049 |
| Residuals | | | | | | | |  | 370963.791 | | | | |  | 28 |  | 13248.707 |  |  |  |  |  |  |
| *Note.*  Type III Sum of Squares | | | | | | | | | | | | | | | | | | | | | | | |
| **Between Subjects Effects** | | | | | | | | | | | | | |  |  |  |  |  |  |  |  |  |  |
| **Cases** | | **Sum of Squares** | | **df** | | **Mean Square** | | **F** | | **p** | | **η²_p_** | |  |  |  |  |  |  |  |  |  |  |
| GROUP |  | 519652.527 |  | 1 |  | 519652.527 |  | 2.847 |  | 0.103 |  | 0.092 |  |  |  |  |  |  |  |  |  |  |  |
| Residuals |  | 5.111×10^+6^ |  | 28 |  | 182527.182 |  |  |  |  |  |  |  |  |  |  |  |  |  |  |  |  |  |
| *Note.*  Type III Sum of Squares | | | | | | | | | | | | | |  |  |  |  |  |  |  |  |  |  |

| **SUPPLEMENTARY TABLE 2b – Grasping Asynchrony - Post Hoc Comparisons - Interaction type ✻ Movement type ✻ Correction** | | | | | | | | | | |
| --- | --- | --- | --- | --- | --- | --- | --- | --- | --- | --- |
|  | |  | | **Mean Difference** | | **SE** | | **t** | | **p_bonf_** |
| Complementary , Precision, Corr |  | Imitative, Precision, Corr |  | 112.303 |  | 22.394 |  | 5.015 |  | < .001 |
|  |  | Complementary , Power, Corr |  | 16.663 |  | 27.024 |  | 0.617 |  | 1.000 |
|  |  | Imitative, Power, Corr |  | -19.020 |  | 24.179 |  | -0.787 |  | 1.000 |
|  |  | Complementary , Precision, NoCorr |  | 141.585 |  | 24.695 |  | 5.733 |  | < .001 |
|  |  | Imitative, Precision, NoCorr |  | 68.005 |  | 21.244 |  | 3.201 |  | 0.051 |
|  |  | Complementary , Power, NoCorr |  | -18.382 |  | 25.954 |  | -0.708 |  | 1.000 |
|  |  | Imitative, Power, NoCorr |  | 132.243 |  | 27.407 |  | 4.825 |  | < .001 |
| Imitative, Precision, Corr |  | Complementary , Power, Corr |  | -95.640 |  | 24.179 |  | -3.956 |  | 0.004 |
|  |  | Imitative, Power, Corr |  | -131.323 |  | 27.024 |  | -4.860 |  | < .001 |
|  |  | Complementary , Precision, NoCorr |  | 29.282 |  | 21.244 |  | 1.378 |  | 1.000 |
|  |  | Imitative, Precision, NoCorr |  | -44.298 |  | 24.695 |  | -1.794 |  | 1.000 |
|  |  | Complementary , Power, NoCorr |  | -130.685 |  | 27.407 |  | -4.768 |  | < .001 |
|  |  | Imitative, Power, NoCorr |  | 19.940 |  | 25.954 |  | 0.768 |  | 1.000 |
| Complementary , Power, Corr |  | Imitative, Power, Corr |  | -35.683 |  | 22.394 |  | -1.593 |  | 1.000 |
|  |  | Complementary , Precision, NoCorr |  | 124.921 |  | 25.954 |  | 4.813 |  | < .001 |
|  |  | Imitative, Precision, NoCorr |  | 51.342 |  | 27.407 |  | 1.873 |  | 1.000 |
|  |  | Complementary , Power, NoCorr |  | -35.045 |  | 24.695 |  | -1.419 |  | 1.000 |
|  |  | Imitative, Power, NoCorr |  | 115.580 |  | 21.244 |  | 5.440 |  | < .001 |
| Imitative, Power, Corr |  | Complementary , Precision, NoCorr |  | 160.604 |  | 27.407 |  | 5.860 |  | < .001 |
|  |  | Imitative, Precision, NoCorr |  | 87.025 |  | 25.954 |  | 3.353 |  | 0.032 |
|  |  | Complementary , Power, NoCorr |  | 0.638 |  | 21.244 |  | 0.030 |  | 1.000 |
|  |  | Imitative, Power, NoCorr |  | 151.263 |  | 24.695 |  | 6.125 |  | < .001 |
| Complementary , Precision, NoCorr |  | Imitative, Precision, NoCorr |  | -73.580 |  | 22.394 |  | -3.286 |  | 0.040 |
|  |  | Complementary , Power, NoCorr |  | -159.966 |  | 27.024 |  | -5.919 |  | < .001 |
|  |  | Imitative, Power, NoCorr |  | -9.342 |  | 24.179 |  | -0.386 |  | 1.000 |
| Imitative, Precision, NoCorr |  | Complementary , Power, NoCorr |  | -86.387 |  | 24.179 |  | -3.573 |  | 0.017 |
|  |  | Imitative, Power, NoCorr |  | 64.238 |  | 27.024 |  | 2.377 |  | 0.545 |
| Complementary , Power, NoCorr |  | Imitative, Power, NoCorr |  | 150.625 |  | 22.394 |  | 6.726 |  | < .001 |
| *Note.*  P-value adjusted for comparing a family of 28 | | | | | | | | | | |
| *Note.*  Results are averaged over the levels of: GROUP | | | | | | | | | | |

| **SUPPLEMENTARY TABLE 3a – Grasping Asynchrony - Mixed ANOVA PD OFF vs HC** | | | | | | | | | | | | | | | | | | | | | | | |
| --- | --- | --- | --- | --- | --- | --- | --- | --- | --- | --- | --- | --- | --- | --- | --- | --- | --- | --- | --- | --- | --- | --- | --- |
| **Within Subjects Effects** | | | | | | | | | | | | | | | | | | | | | | | |
| **Cases** | | | | | | | | | | | **Sum of Squares** | | | | **df** | | **Mean Square** | | **F** | | **p** | | **η²_p_** |
| Interaction type | | | | | | | | | |  | 78938.167 | | |  | 1 |  | 78938.167 |  | 15.107 |  | < .001 |  | 0.350 |
| Interaction type ✻ GROUP | | | | | | | | | |  | 2955.286 | | |  | 1 |  | 2955.286 |  | 0.566 |  | 0.458 |  | 0.020 |
| Residuals | | | | | | | | | |  | 146310.013 | | |  | 28 |  | 5225.358 |  |  |  |  |  |  |
| Movement type | | | | | | | | | |  | 196123.706 | | |  | 1 |  | 196123.706 |  | 12.834 |  | 0.001 |  | 0.314 |
| Movement type ✻ GROUP | | | | | | | | | |  | 497.817 | | |  | 1 |  | 497.817 |  | 0.033 |  | 0.858 |  | 0.001 |
| Residuals | | | | | | | | | |  | 427879.923 | | |  | 28 |  | 15281.426 |  |  |  |  |  |  |
| Correction | | | | | | | | | |  | 241226.340 | | |  | 1 |  | 241226.340 |  | 25.682 |  | < .001 |  | 0.478 |
| Correction ✻ GROUP | | | | | | | | | |  | 156.647 | | |  | 1 |  | 156.647 |  | 0.017 |  | 0.898 |  | 5.953×10^-4^ |
| Residuals | | | | | | | | | |  | 262996.209 | | |  | 28 |  | 9392.722 |  |  |  |  |  |  |
| Interaction type ✻ Movement type | | | | | | | | | |  | 14573.324 | | |  | 1 |  | 14573.324 |  | 2.470 |  | 0.127 |  | 0.081 |
| Interaction type ✻ Movement type ✻ GROUP | | | | | | | | | |  | 220.211 | | |  | 1 |  | 220.211 |  | 0.037 |  | 0.848 |  | 0.001 |
| Residuals | | | | | | | | | |  | 165237.109 | | |  | 28 |  | 5901.325 |  |  |  |  |  |  |
| Interaction type ✻ Correction | | | | | | | | | |  | 8.592 | | |  | 1 |  | 8.592 |  | 0.001 |  | 0.970 |  | 5.036×10^-5^ |
| Interaction type ✻ Correction ✻ GROUP | | | | | | | | | |  | 9144.233 | | |  | 1 |  | 9144.233 |  | 1.501 |  | 0.231 |  | 0.051 |
| Residuals | | | | | | | | | |  | 170603.565 | | |  | 28 |  | 6092.984 |  |  |  |  |  |  |
| Movement type ✻ Correction | | | | | | | | | |  | 6082.987 | | |  | 1 |  | 6082.987 |  | 1.403 |  | 0.246 |  | 0.048 |
| Movement type ✻ Correction ✻ GROUP | | | | | | | | | |  | 7793.984 | | |  | 1 |  | 7793.984 |  | 1.798 |  | 0.191 |  | 0.060 |
| Residuals | | | | | | | | | |  | 121378.173 | | |  | 28 |  | 4334.935 |  |  |  |  |  |  |
| Interaction type ✻ Movement type ✻ Correction | | | | | | | | | |  | 421106.648 | | |  | 1 |  | 421106.648 |  | 26.238 |  | < .001 |  | 0.484 |
| Interaction type ✻ Movement type ✻ Correction ✻ GROUP | | | | | | | | | |  | 44246.813 | | |  | 1 |  | 44246.813 |  | 2.757 |  | 0.108 |  | 0.090 |
| Residuals | | | | | | | | | |  | 449385.369 | | |  | 28 |  | 16049.477 |  |  |  |  |  |  |
| *Note.*  Type III Sum of Squares | | | | | | | | | | | | | | | | | | | | | | | |
| **Between Subjects Effects** | | | | | | | | | | | | | |  |  |  |  |  |  |  |  |  |  |
| **Cases** | | **Sum of Squares** | | **df** | | **Mean Square** | | **F** | | **p** | | **η²_p_** | |  |  |  |  |  |  |  |  |  |  |
| GROUP |  | 1.668×10^+6^ |  | 1 |  | 1.668×10^+6^ |  | 7.384 |  | 0.011 |  | 0.209 |  |  |  |  |  |  |  |  |  |  |  |
| Residuals |  | 6.324×10^+6^ |  | 28 |  | 225868.228 |  |  |  |  |  |  |  |  |  |  |  |  |  |  |  |  |  |
| *Note.*  Type III Sum of Squares | | | | | | | | | | | | | |  |  |  |  |  |  |  |  |  |  |

| **SUPPLEMENTARY TABLE 3b – Grasping Asynchrony - Post Hoc Comparisons - Interaction type ✻ Movement type ✻ Correction** | | | | | | | | | | |
| --- | --- | --- | --- | --- | --- | --- | --- | --- | --- | --- |
|  | |  | | **Mean Difference** | | **SE** | | **t** | | **p_bonf_** |
| Complementary, Precision, Corr |  | Imitative, Precision, Corr |  | 104.085 |  | 23.548 |  | 4.420 |  | < .001 |
|  |  | Complementary, Power, Corr |  | 0.950 |  | 26.321 |  | 0.036 |  | 1.000 |
|  |  | Imitative, Power, Corr |  | -31.348 |  | 22.706 |  | -1.381 |  | 1.000 |
|  |  | Complementary, Precision, NoCorr |  | 136.736 |  | 24.451 |  | 5.592 |  | < .001 |
|  |  | Imitative, Precision, NoCorr |  | 74.025 |  | 20.353 |  | 3.637 |  | 0.012 |
|  |  | Complementary, Power, NoCorr |  | -9.729 |  | 24.721 |  | -0.394 |  | 1.000 |
|  |  | Imitative, Power, NoCorr |  | 126.282 |  | 27.673 |  | 4.563 |  | < .001 |
| Imitative, Precision, Corr |  | Complementary, Power, Corr |  | -103.135 |  | 22.706 |  | -4.542 |  | < .001 |
|  |  | Imitative, Power, Corr |  | -135.433 |  | 26.321 |  | -5.145 |  | < .001 |
|  |  | Complementary, Precision, NoCorr |  | 32.651 |  | 20.353 |  | 1.604 |  | 1.000 |
|  |  | Imitative, Precision, NoCorr |  | -30.060 |  | 24.451 |  | -1.229 |  | 1.000 |
|  |  | Complementary, Power, NoCorr |  | -113.814 |  | 27.673 |  | -4.113 |  | 0.002 |
|  |  | Imitative, Power, NoCorr |  | 22.197 |  | 24.721 |  | 0.898 |  | 1.000 |
| Complementary, Power, Corr |  | Imitative, Power, Corr |  | -32.298 |  | 23.548 |  | -1.372 |  | 1.000 |
|  |  | Complementary, Precision, NoCorr |  | 135.786 |  | 24.721 |  | 5.493 |  | < .001 |
|  |  | Imitative, Precision, NoCorr |  | 73.075 |  | 27.673 |  | 2.641 |  | 0.270 |
|  |  | Complementary, Power, NoCorr |  | -10.679 |  | 24.451 |  | -0.437 |  | 1.000 |
|  |  | Imitative, Power, NoCorr |  | 125.332 |  | 20.353 |  | 6.158 |  | < .001 |
| Imitative, Power, Corr |  | Complementary, Precision, NoCorr |  | 168.084 |  | 27.673 |  | 6.074 |  | < .001 |
|  |  | Imitative, Precision, NoCorr |  | 105.373 |  | 24.721 |  | 4.262 |  | 0.001 |
|  |  | Complementary, Power, NoCorr |  | 21.619 |  | 20.353 |  | 1.062 |  | 1.000 |
|  |  | Imitative, Power, NoCorr |  | 157.630 |  | 24.451 |  | 6.447 |  | < .001 |
| Complementary, Precision, NoCorr |  | Imitative, Precision, NoCorr |  | -62.711 |  | 23.548 |  | -2.663 |  | 0.258 |
|  |  | Complementary, Power, NoCorr |  | -146.465 |  | 26.321 |  | -5.565 |  | < .001 |
|  |  | Imitative, Power, NoCorr |  | -10.454 |  | 22.706 |  | -0.460 |  | 1.000 |
| Imitative, Precision, NoCorr |  | Complementary, Power, NoCorr |  | -83.754 |  | 22.706 |  | -3.689 |  | 0.011 |
|  |  | Imitative, Power, NoCorr |  | 52.257 |  | 26.321 |  | 1.985 |  | 1.000 |
| Complementary, Power, NoCorr |  | Imitative, Power, NoCorr |  | 136.011 |  | 23.548 |  | 5.776 |  | < .001 |
| *Note.*  P-value adjusted for comparing a family of 28 | | | | | | | | | | |
| *Note.*  Results are averaged over the levels of: GROUP | | | | | | | | | | |

| **SUPPLEMENTARY TABLE 4a – Movement time - Within-participants ANOVA PD ON vs PD OFF** | | | | | | | | | | | | |
| --- | --- | --- | --- | --- | --- | --- | --- | --- | --- | --- | --- | --- |
| **Within Subjects Effects** | | | | | | | | | | | | |
| **Cases** | | **Sum of Squares** | | **df** | | **Mean Square** | | **F** | | **p** | | **η²_p_** |
| Condition |  | 31974.232 |  | 1 |  | 31974.232 |  | 0.273 |  | 0.609 |  | 0.019 |
| Residuals |  | 1.640×10^+6^ |  | 14 |  | 117120.483 |  |  |  |  |  |  |
| Correction |  | 53329.783 |  | 1 |  | 53329.783 |  | 2.609 |  | 0.129 |  | 0.157 |
| Residuals |  | 286172.819 |  | 14 |  | 20440.916 |  |  |  |  |  |  |
| Interaction type |  | 223800.675 |  | 1 |  | 223800.675 |  | 20.006 |  | < .001 |  | 0.588 |
| Residuals |  | 156610.873 |  | 14 |  | 11186.491 |  |  |  |  |  |  |
| Movement type |  | 1057.558 |  | 1 |  | 1057.558 |  | 0.038 |  | 0.848 |  | 0.003 |
| Residuals |  | 386261.834 |  | 14 |  | 27590.131 |  |  |  |  |  |  |
| Condition ✻ Correction |  | 634.065 |  | 1 |  | 634.065 |  | 0.149 |  | 0.706 |  | 0.011 |
| Residuals |  | 59696.705 |  | 14 |  | 4264.050 |  |  |  |  |  |  |
| Condition ✻ Interaction type |  | 4461.432 |  | 1 |  | 4461.432 |  | 0.389 |  | 0.543 |  | 0.027 |
| Residuals |  | 160686.783 |  | 14 |  | 11477.627 |  |  |  |  |  |  |
| Correction ✻ Interaction type |  | 1170.630 |  | 1 |  | 1170.630 |  | 0.310 |  | 0.586 |  | 0.022 |
| Residuals |  | 52845.032 |  | 14 |  | 3774.645 |  |  |  |  |  |  |
| Condition ✻ Movement type |  | 14712.979 |  | 1 |  | 14712.979 |  | 1.204 |  | 0.291 |  | 0.079 |
| Residuals |  | 171016.589 |  | 14 |  | 12215.471 |  |  |  |  |  |  |
| Correction ✻ Movement type |  | 4751.655 |  | 1 |  | 4751.655 |  | 0.498 |  | 0.492 |  | 0.034 |
| Residuals |  | 133568.762 |  | 14 |  | 9540.626 |  |  |  |  |  |  |
| Interaction type ✻ Movement type |  | 21404.016 |  | 1 |  | 21404.016 |  | 1.101 |  | 0.312 |  | 0.073 |
| Residuals |  | 272222.756 |  | 14 |  | 19444.483 |  |  |  |  |  |  |
| Condition ✻ Correction ✻ Interaction type |  | 5721.873 |  | 1 |  | 5721.873 |  | 0.744 |  | 0.403 |  | 0.050 |
| Residuals |  | 107699.349 |  | 14 |  | 7692.811 |  |  |  |  |  |  |
| Condition ✻ Correction ✻ Movement type |  | 96.668 |  | 1 |  | 96.668 |  | 0.015 |  | 0.904 |  | 0.001 |
| Residuals |  | 90321.587 |  | 14 |  | 6451.542 |  |  |  |  |  |  |
| Condition ✻ Interaction type ✻ Movement type |  | 37298.655 |  | 1 |  | 37298.655 |  | 4.212 |  | 0.059 |  | 0.231 |
| Residuals |  | 123972.830 |  | 14 |  | 8855.202 |  |  |  |  |  |  |
| Correction ✻ Interaction type ✻ Movement type |  | 124864.533 |  | 1 |  | 124864.533 |  | 9.861 |  | 0.007 |  | 0.413 |
| Residuals |  | 177280.933 |  | 14 |  | 12662.924 |  |  |  |  |  |  |
| Condition ✻ Correction ✻ Interaction type ✻ Movement type |  | 908.691 |  | 1 |  | 908.691 |  | 0.246 |  | 0.627 |  | 0.017 |
| Residuals |  | 51642.319 |  | 14 |  | 3688.737 |  |  |  |  |  |  |
| *Note.*  Type III Sum of Squares | | | | | | | | | | | | |

| **SUPPLEMENTARY TABLE 4b – Movement time - Post Hoc Comparisons - Correction ✻ Interaction type ✻ Movement type** | | | | | | | | | | |
| --- | --- | --- | --- | --- | --- | --- | --- | --- | --- | --- |
|  | |  | | **Mean Difference** | | **SE** | | **t** | | **p_bonf_** |
| Corr, Complementary, Precision |  | NoCorr, Complementary, Precision |  | -2.489 |  | 27.815 |  | -0.089 |  | 1.000 |
|  |  | Corr, Imitative, Precision |  | 0.985 |  | 28.008 |  | 0.035 |  | 1.000 |
|  |  | NoCorr, Imitative, Precision |  | 80.899 |  | 31.784 |  | 2.545 |  | 0.392 |
|  |  | Corr, Complementary, Power |  | -59.805 |  | 33.970 |  | -1.761 |  | 1.000 |
|  |  | NoCorr, Complementary, Power |  | 11.145 |  | 34.460 |  | 0.323 |  | 1.000 |
|  |  | Corr, Imitative, Power |  | 70.192 |  | 29.465 |  | 2.382 |  | 0.624 |
|  |  | NoCorr, Imitative, Power |  | 41.070 |  | 34.612 |  | 1.187 |  | 1.000 |
| NoCorr, Complementary, Precision |  | Corr, Imitative, Precision |  | 3.474 |  | 31.784 |  | 0.109 |  | 1.000 |
|  |  | NoCorr, Imitative, Precision |  | 83.388 |  | 28.008 |  | 2.977 |  | 0.130 |
|  |  | Corr, Complementary, Power |  | -57.316 |  | 34.460 |  | -1.663 |  | 1.000 |
|  |  | NoCorr, Complementary, Power |  | 13.634 |  | 33.970 |  | 0.401 |  | 1.000 |
|  |  | Corr, Imitative, Power |  | 72.681 |  | 34.612 |  | 2.100 |  | 1.000 |
|  |  | NoCorr, Imitative, Power |  | 43.559 |  | 29.465 |  | 1.478 |  | 1.000 |
| Corr, Imitative, Precision |  | NoCorr, Imitative, Precision |  | 79.914 |  | 27.815 |  | 2.873 |  | 0.174 |
|  |  | Corr, Complementary, Power |  | -60.790 |  | 29.465 |  | -2.063 |  | 1.000 |
|  |  | NoCorr, Complementary, Power |  | 10.160 |  | 34.612 |  | 0.294 |  | 1.000 |
|  |  | Corr, Imitative, Power |  | 69.207 |  | 33.970 |  | 2.037 |  | 1.000 |
|  |  | NoCorr, Imitative, Power |  | 40.085 |  | 34.460 |  | 1.163 |  | 1.000 |
| NoCorr, Imitative, Precision |  | Corr, Complementary, Power |  | -140.704 |  | 34.612 |  | -4.065 |  | 0.005 |
|  |  | NoCorr, Complementary, Power |  | -69.754 |  | 29.465 |  | -2.367 |  | 0.646 |
|  |  | Corr, Imitative, Power |  | -10.707 |  | 34.460 |  | -0.311 |  | 1.000 |
|  |  | NoCorr, Imitative, Power |  | -39.829 |  | 33.970 |  | -1.172 |  | 1.000 |
| Corr, Complementary, Power |  | NoCorr, Complementary, Power |  | 70.950 |  | 27.815 |  | 2.551 |  | 0.400 |
|  |  | Corr, Imitative, Power |  | 129.997 |  | 28.008 |  | 4.641 |  | < .001 |
|  |  | NoCorr, Imitative, Power |  | 100.875 |  | 31.784 |  | 3.174 |  | 0.072 |
| NoCorr, Complementary, Power |  | Corr, Imitative, Power |  | 59.047 |  | 31.784 |  | 1.858 |  | 1.000 |
|  |  | NoCorr, Imitative, Power |  | 29.925 |  | 28.008 |  | 1.068 |  | 1.000 |
| Corr, Imitative, Power |  | NoCorr, Imitative, Power |  | -29.122 |  | 27.815 |  | -1.047 |  | 1.000 |
| *Note.*  P-value adjusted for comparing a family of 28 | | | | | | | | | | |
| *Note.*  Results are averaged over the levels of: Condition | | | | | | | | | | |

| **SUPPLEMENTARY TABLE 5a – Movement Time - Mixed ANOVA PD ON vs HC** | | | | | | | | | | | | |
| --- | --- | --- | --- | --- | --- | --- | --- | --- | --- | --- | --- | --- |
| **Within Subjects Effects** | | | | | | | | | | | | |
| **Cases** | | **Sum of Squares** | | **df** | | **Mean Square** | | **F** | | **p** | | **η²_p_** |
| Correction |  | 59644.001 |  | 1 |  | 59644.001 |  | 4.223 |  | 0.049 |  | 0.131 |
| Correction ✻ GROUP |  | 1479.897 |  | 1 |  | 1479.897 |  | 0.105 |  | 0.749 |  | 0.004 |
| Residuals |  | 395451.054 |  | 28 |  | 14123.252 |  |  |  |  |  |  |
| Interaction type |  | 137407.526 |  | 1 |  | 137407.526 |  | 14.174 |  | < .001 |  | 0.336 |
| Interaction type ✻ GROUP |  | 28623.396 |  | 1 |  | 28623.396 |  | 2.953 |  | 0.097 |  | 0.095 |
| Residuals |  | 271438.209 |  | 28 |  | 9694.222 |  |  |  |  |  |  |
| Movement type |  | 11831.106 |  | 1 |  | 11831.106 |  | 0.963 |  | 0.335 |  | 0.033 |
| Movement type ✻ GROUP |  | 399.757 |  | 1 |  | 399.757 |  | 0.033 |  | 0.858 |  | 0.001 |
| Residuals |  | 344157.343 |  | 28 |  | 12291.334 |  |  |  |  |  |  |
| Correction ✻ Interaction type |  | 2512.430 |  | 1 |  | 2512.430 |  | 0.404 |  | 0.530 |  | 0.014 |
| Correction ✻ Interaction type ✻ GROUP |  | 3568.078 |  | 1 |  | 3568.078 |  | 0.574 |  | 0.455 |  | 0.020 |
| Residuals |  | 174100.615 |  | 28 |  | 6217.879 |  |  |  |  |  |  |
| Correction ✻ Movement type |  | 16647.268 |  | 1 |  | 16647.268 |  | 1.999 |  | 0.168 |  | 0.067 |
| Correction ✻ Movement type ✻ GROUP |  | 2526.071 |  | 1 |  | 2526.071 |  | 0.303 |  | 0.586 |  | 0.011 |
| Residuals |  | 233224.395 |  | 28 |  | 8329.443 |  |  |  |  |  |  |
| Interaction type ✻ Movement type |  | 20509.543 |  | 1 |  | 20509.543 |  | 1.307 |  | 0.263 |  | 0.045 |
| Interaction type ✻ Movement type ✻ GROUP |  | 38501.574 |  | 1 |  | 38501.574 |  | 2.454 |  | 0.128 |  | 0.081 |
| Residuals |  | 439261.787 |  | 28 |  | 15687.921 |  |  |  |  |  |  |
| Correction ✻ Interaction type ✻ Movement type |  | 40306.091 |  | 1 |  | 40306.091 |  | 5.501 |  | 0.026 |  | 0.164 |
| Correction ✻ Interaction type ✻ Movement type ✻ GROUP |  | 14994.859 |  | 1 |  | 14994.859 |  | 2.046 |  | 0.164 |  | 0.068 |
| Residuals |  | 205171.024 |  | 28 |  | 7327.537 |  |  |  |  |  |  |
| *Note.*  Type III Sum of Squares | | | | | | | | | | | | |

| **Between Subjects Effects** | | | | | | | | | | | | |
| --- | --- | --- | --- | --- | --- | --- | --- | --- | --- | --- | --- | --- |
| **Cases** | | **Sum of Squares** | | **df** | | **Mean Square** | | **F** | | **p** | | **η²_p_** |
| GROUP |  | 88076.752 |  | 1 |  | 88076.752 |  | 0.189 |  | 0.667 |  | 0.007 |
| Residuals |  | 1.308×10^+7^ |  | 28 |  | 467061.669 |  |  |  |  |  |  |
| *Note.*  Type III Sum of Squares | | | | | | | | | | | | |

**SUPPLEMENTARY TABLE 6a – Movement Time - Mixed ANOVA PD OFF vs HC**

|  | | | | | | | | | | | | |
| --- | --- | --- | --- | --- | --- | --- | --- | --- | --- | --- | --- | --- |
| **Within Subjects Effects** | | | | | | | | | | | | |
| **Cases** | | **Sum of Squares** | | **df** | | **Mean Square** | | **F** | | **p** | | **η²_p_** |
| Correction |  | 72577.360 |  | 1 |  | 72577.360 |  | 6.069 |  | 0.020 |  | 0.178 |
| Correction ✻ GROUP |  | 176.592 |  | 1 |  | 176.592 |  | 0.015 |  | 0.904 |  | 5.271×10^-4^ |
| Residuals |  | 334820.342 |  | 28 |  | 11957.869 |  |  |  |  |  |  |
| Interaction type |  | 92349.897 |  | 1 |  | 92349.897 |  | 10.266 |  | 0.003 |  | 0.268 |
| Interaction type ✻ GROUP |  | 10483.825 |  | 1 |  | 10483.825 |  | 1.165 |  | 0.290 |  | 0.040 |
| Residuals |  | 251875.593 |  | 28 |  | 8995.557 |  |  |  |  |  |  |
| Movement type |  | 156.905 |  | 1 |  | 156.905 |  | 0.009 |  | 0.926 |  | 3.137×10^-4^ |
| Movement type ✻ GROUP |  | 19963.146 |  | 1 |  | 19963.146 |  | 1.118 |  | 0.299 |  | 0.038 |
| Residuals |  | 500073.242 |  | 28 |  | 17859.759 |  |  |  |  |  |  |
| Correction ✻ Interaction type |  | 651.215 |  | 1 |  | 651.215 |  | 0.095 |  | 0.760 |  | 0.003 |
| Correction ✻ Interaction type ✻ GROUP |  | 253.118 |  | 1 |  | 253.118 |  | 0.037 |  | 0.849 |  | 0.001 |
| Residuals |  | 191197.916 |  | 28 |  | 6828.497 |  |  |  |  |  |  |
| Correction ✻ Movement type |  | 14206.801 |  | 1 |  | 14206.801 |  | 4.226 |  | 0.049 |  | 0.131 |
| Correction ✻ Movement type ✻ GROUP |  | 3611.053 |  | 1 |  | 3611.053 |  | 1.074 |  | 0.309 |  | 0.037 |
| Residuals |  | 94125.850 |  | 28 |  | 3361.638 |  |  |  |  |  |  |
| Interaction type ✻ Movement type |  | 2491.714 |  | 1 |  | 2491.714 |  | 0.300 |  | 0.588 |  | 0.011 |
| Interaction type ✻ Movement type ✻ GROUP |  | 9.546 |  | 1 |  | 9.546 |  | 0.001 |  | 0.973 |  | 4.100×10^-5^ |
| Residuals |  | 232783.467 |  | 28 |  | 8313.695 |  |  |  |  |  |  |
| Correction ✻ Interaction type ✻ Movement type |  | 53318.625 |  | 1 |  | 53318.625 |  | 7.938 |  | 0.009 |  | 0.221 |
| Correction ✻ Interaction type ✻ Movement type ✻ GROUP |  | 23286.147 |  | 1 |  | 23286.147 |  | 3.467 |  | 0.073 |  | 0.110 |
| Residuals |  | 188067.698 |  | 28 |  | 6716.703 |  |  |  |  |  |  |
| *Note.*  Type III Sum of Squares | | | | | | | | | | | | |

| **Between Subjects Effects** | | | | | | | | | | | | |
| --- | --- | --- | --- | --- | --- | --- | --- | --- | --- | --- | --- | --- |
| **Cases** | | **Sum of Squares** | | **df** | | **Mean Square** | | **F** | | **p** | | **η²_p_** |
| GROUP |  | 226186.493 |  | 1 |  | 226186.493 |  | 0.630 |  | 0.434 |  | 0.022 |
| Residuals |  | 1.005×10^+7^ |  | 28 |  | 358926.304 |  |  |  |  |  |  |
| *Note.*  Type III Sum of Squares | | | | | | | | | | | | |

| **SUPPLEMENTARY TABLE 7a – Reaction Times - Within-participants ANOVA PD ON vs PD OFF** | | | | | | | | | | | | |
| --- | --- | --- | --- | --- | --- | --- | --- | --- | --- | --- | --- | --- |
| **Within Subjects Effects** | | | | | | | | | | | | |
| **Cases** | | **Sum of Squares** | | **df** | | **Mean Square** | | **F** | | **p** | | **η²_p_** |
| Condition |  | 71590.025 |  | 1 |  | 71590.025 |  | 0.217 |  | 0.648 |  | 0.015 |
| Residuals |  | 4.616×10^+6^ |  | 14 |  | 329723.326 |  |  |  |  |  |  |
| Correction |  | 30464.100 |  | 1 |  | 30464.100 |  | 0.893 |  | 0.361 |  | 0.060 |
| Residuals |  | 477648.047 |  | 14 |  | 34117.718 |  |  |  |  |  |  |
| Interaction type |  | 50765.450 |  | 1 |  | 50765.450 |  | 0.777 |  | 0.393 |  | 0.053 |
| Residuals |  | 914660.018 |  | 14 |  | 65332.858 |  |  |  |  |  |  |
| Movement type |  | 56850.114 |  | 1 |  | 56850.114 |  | 4.186 |  | 0.060 |  | 0.230 |
| Residuals |  | 190139.796 |  | 14 |  | 13581.414 |  |  |  |  |  |  |
| Condition ✻ Correction |  | 78890.572 |  | 1 |  | 78890.572 |  | 1.114 |  | 0.309 |  | 0.074 |
| Residuals |  | 991436.158 |  | 14 |  | 70816.868 |  |  |  |  |  |  |
| Condition ✻ Interaction type |  | 84564.673 |  | 1 |  | 84564.673 |  | 2.622 |  | 0.128 |  | 0.158 |
| Residuals |  | 451549.220 |  | 14 |  | 32253.516 |  |  |  |  |  |  |
| Correction ✻ Interaction type |  | 38249.862 |  | 1 |  | 38249.862 |  | 1.032 |  | 0.327 |  | 0.069 |
| Residuals |  | 519101.189 |  | 14 |  | 37078.656 |  |  |  |  |  |  |
| Condition ✻ Movement type |  | 55892.193 |  | 1 |  | 55892.193 |  | 2.999 |  | 0.105 |  | 0.176 |
| Residuals |  | 260895.446 |  | 14 |  | 18635.389 |  |  |  |  |  |  |
| Correction ✻ Movement type |  | 3149.180 |  | 1 |  | 3149.180 |  | 0.169 |  | 0.688 |  | 0.012 |
| Residuals |  | 261411.231 |  | 14 |  | 18672.231 |  |  |  |  |  |  |
| Interaction type ✻ Movement type |  | 18116.739 |  | 1 |  | 18116.739 |  | 3.332 |  | 0.089 |  | 0.192 |
| Residuals |  | 76117.187 |  | 14 |  | 5436.942 |  |  |  |  |  |  |
| Condition ✻ Correction ✻ Interaction type |  | 2135.906 |  | 1 |  | 2135.906 |  | 0.053 |  | 0.821 |  | 0.004 |
| Residuals |  | 563511.703 |  | 14 |  | 40250.836 |  |  |  |  |  |  |
| Condition ✻ Correction ✻ Movement type |  | 2448.870 |  | 1 |  | 2448.870 |  | 0.114 |  | 0.741 |  | 0.008 |
| Residuals |  | 300511.251 |  | 14 |  | 21465.089 |  |  |  |  |  |  |
| Condition ✻ Interaction type ✻ Movement type |  | 4.928 |  | 1 |  | 4.928 |  | 4.994×10^-4^ |  | 0.982 |  | 3.567×10^-5^ |
| Residuals |  | 138144.071 |  | 14 |  | 9867.434 |  |  |  |  |  |  |
| Correction ✻ Interaction type ✻ Movement type |  | 25141.704 |  | 1 |  | 25141.704 |  | 1.555 |  | 0.233 |  | 0.100 |
| Residuals |  | 226283.651 |  | 14 |  | 16163.118 |  |  |  |  |  |  |
| Condition ✻ Correction ✻ Interaction type ✻ Movement type |  | 930.734 |  | 1 |  | 930.734 |  | 0.036 |  | 0.852 |  | 0.003 |
| Residuals |  | 360205.054 |  | 14 |  | 25728.932 |  |  |  |  |  |  |
| *Note.*  Type III Sum of Squares | | | | | | | | | | | | |

| **SUPPLEMENTARY TABLE 8a – Reaction Times - Mixed ANOVA PD ON vs HC** | | | | | | | | | | | | |
| --- | --- | --- | --- | --- | --- | --- | --- | --- | --- | --- | --- | --- |
| **Within Subjects Effects** | | | | | | | | | | | | |
| **Cases** | | **Sum of Squares** | | **df** | | **Mean Square** | | **F** | | **p** | | **η²_p_** |
| Correction |  | 270372.770 |  | 1 |  | 270372.770 |  | 3.417 |  | 0.075 |  | 0.109 |
| Correction ✻ GROUP |  | 4167.927 |  | 1 |  | 4167.927 |  | 0.053 |  | 0.820 |  | 0.002 |
| Residuals |  | 2.216×10^+6^ |  | 28 |  | 79125.795 |  |  |  |  |  |  |
| Interaction type |  | 56936.107 |  | 1 |  | 56936.107 |  | 1.346 |  | 0.256 |  | 0.046 |
| Interaction type ✻ GROUP |  | 77005.734 |  | 1 |  | 77005.734 |  | 1.821 |  | 0.188 |  | 0.061 |
| Residuals |  | 1.184×10^+6^ |  | 28 |  | 42296.465 |  |  |  |  |  |  |
| Movement type |  | 89457.064 |  | 1 |  | 89457.064 |  | 8.898 |  | 0.006 |  | 0.241 |
| Movement type ✻ GROUP |  | 30889.531 |  | 1 |  | 30889.531 |  | 3.073 |  | 0.091 |  | 0.099 |
| Residuals |  | 281487.667 |  | 28 |  | 10053.131 |  |  |  |  |  |  |
| Correction ✻ Interaction type |  | 22183.205 |  | 1 |  | 22183.205 |  | 0.610 |  | 0.441 |  | 0.021 |
| Correction ✻ Interaction type ✻ GROUP |  | 8621.367 |  | 1 |  | 8621.367 |  | 0.237 |  | 0.630 |  | 0.008 |
| Residuals |  | 1.018×10^+6^ |  | 28 |  | 36340.797 |  |  |  |  |  |  |
| Correction ✻ Movement type |  | 6901.388 |  | 1 |  | 6901.388 |  | 0.354 |  | 0.557 |  | 0.012 |
| Correction ✻ Movement type ✻ GROUP |  | 507.557 |  | 1 |  | 507.557 |  | 0.026 |  | 0.873 |  | 9.282×10^-4^ |
| Residuals |  | 546302.510 |  | 28 |  | 19510.804 |  |  |  |  |  |  |
| Interaction type ✻ Movement type |  | 4275.664 |  | 1 |  | 4275.664 |  | 0.321 |  | 0.576 |  | 0.011 |
| Interaction type ✻ Movement type ✻ GROUP |  | 5102.210 |  | 1 |  | 5102.210 |  | 0.383 |  | 0.541 |  | 0.013 |
| Residuals |  | 373261.341 |  | 28 |  | 13330.762 |  |  |  |  |  |  |
| Correction ✻ Interaction type ✻ Movement type |  | 31348.072 |  | 1 |  | 31348.072 |  | 1.230 |  | 0.277 |  | 0.042 |
| Correction ✻ Interaction type ✻ Movement type ✻ GROUP |  | 144.370 |  | 1 |  | 144.370 |  | 0.006 |  | 0.941 |  | 2.023×10^-4^ |
| Residuals |  | 713622.307 |  | 28 |  | 25486.511 |  |  |  |  |  |  |
| *Note.*  Type III Sum of Squares | | | | | | | | | | | | |

| **Between Subjects Effects** | | | | | | | | | | | | |
| --- | --- | --- | --- | --- | --- | --- | --- | --- | --- | --- | --- | --- |
| **Cases** | | **Sum of Squares** | | **df** | | **Mean Square** | | **F** | | **p** | | **η²_p_** |
| GROUP |  | 63830.325 |  | 1 |  | 63830.325 |  | 0.168 |  | 0.685 |  | 0.006 |
| Residuals |  | 1.063×10^+7^ |  | 28 |  | 379749.959 |  |  |  |  |  |  |
| *Note.*  Type III Sum of Squares | | | | | | | | | | | | |

| **SUPPLEMENTARY TABLE 9a – Reaction Times - Mixed ANOVA PD OFF vs HC** | | | | | | | | | | | | |
| --- | --- | --- | --- | --- | --- | --- | --- | --- | --- | --- | --- | --- |
| **Within Subjects Effects** | | | | | | | | | | | | |
| **Cases** | | **Sum of Squares** | | **df** | | **Mean Square** | | **F** | | **p** | | **η²_p_** |
| Correction |  | 57168.409 |  | 1 |  | 57168.409 |  | 1.350 |  | 0.255 |  | 0.046 |
| Correction ✻ GROUP |  | 119324.743 |  | 1 |  | 119324.743 |  | 2.818 |  | 0.104 |  | 0.091 |
| Residuals |  | 1.185×10^+6^ |  | 28 |  | 42338.444 |  |  |  |  |  |  |
| Interaction type |  | 2723.501 |  | 1 |  | 2723.501 |  | 0.195 |  | 0.662 |  | 0.007 |
| Interaction type ✻ GROUP |  | 176.916 |  | 1 |  | 176.916 |  | 0.013 |  | 0.911 |  | 4.531×10^-4^ |
| Residuals |  | 390268.816 |  | 28 |  | 13938.172 |  |  |  |  |  |  |
| Movement type |  | 3928.587 |  | 1 |  | 3928.587 |  | 0.296 |  | 0.591 |  | 0.010 |
| Movement type ✻ GROUP |  | 3679.771 |  | 1 |  | 3679.771 |  | 0.277 |  | 0.603 |  | 0.010 |
| Residuals |  | 372176.832 |  | 28 |  | 13292.030 |  |  |  |  |  |  |
| Correction ✻ Interaction type |  | 10552.303 |  | 1 |  | 10552.303 |  | 0.737 |  | 0.398 |  | 0.026 |
| Correction ✻ Interaction type ✻ GROUP |  | 2174.865 |  | 1 |  | 2174.865 |  | 0.152 |  | 0.700 |  | 0.005 |
| Residuals |  | 401016.356 |  | 28 |  | 14322.013 |  |  |  |  |  |  |
| Correction ✻ Movement type |  | 1128.190 |  | 1 |  | 1128.190 |  | 0.048 |  | 0.828 |  | 0.002 |
| Correction ✻ Movement type ✻ GROUP |  | 726.682 |  | 1 |  | 726.682 |  | 0.031 |  | 0.862 |  | 0.001 |
| Residuals |  | 656977.993 |  | 28 |  | 23463.500 |  |  |  |  |  |  |
| Interaction type ✻ Movement type |  | 3990.282 |  | 1 |  | 3990.282 |  | 0.295 |  | 0.591 |  | 0.010 |
| Interaction type ✻ Movement type ✻ GROUP |  | 4790.007 |  | 1 |  | 4790.007 |  | 0.354 |  | 0.557 |  | 0.012 |
| Residuals |  | 378937.741 |  | 28 |  | 13533.491 |  |  |  |  |  |  |
| Correction ✻ Interaction type ✻ Movement type |  | 21475.713 |  | 1 |  | 21475.713 |  | 0.861 |  | 0.361 |  | 0.030 |
| Correction ✻ Interaction type ✻ Movement type ✻ GROUP |  | 341.973 |  | 1 |  | 341.973 |  | 0.014 |  | 0.908 |  | 4.892×10^-4^ |
| Residuals |  | 698704.357 |  | 28 |  | 24953.727 |  |  |  |  |  |  |
| *Note.*  Type III Sum of Squares | | | | | | | | | | | | |

| **Between Subjects Effects** | | | | | | | | | | | | |
| --- | --- | --- | --- | --- | --- | --- | --- | --- | --- | --- | --- | --- |
| **Cases** | | **Sum of Squares** | | **df** | | **Mean Square** | | **F** | | **p** | | **η²_p_** |
| GROUP |  | 222.501 |  | 1 |  | 222.501 |  | 8.520×10^-4^ |  | 0.977 |  | 3.043×10^-5^ |
| Residuals |  | 7.313×10^+6^ |  | 28 |  | 261162.717 |  |  |  |  |  |  |
| *Note.*  Type III Sum of Squares | | | | | | | | | | | | |

| **SUPPLEMENTARY TABLE 10 – Pe amplitude - ANOVA PD ON vs OFF** | | | | | | | | | | | | |
| --- | --- | --- | --- | --- | --- | --- | --- | --- | --- | --- | --- | --- |
| **Within Subjects Effects** | | | | | | | | | | | | |
| **Cases** | | **Sum of Squares** | | **df** | | **Mean Square** | | **F** | | **p** | | **η²_p_** |
| Condition |  | 0.009 |  | 1 |  | 0.009 |  | 0.128 |  | 0.725 |  | 0.009 |
| Residuals |  | 1.022 |  | 14 |  | 0.073 |  |  |  |  |  |  |
| Correction |  | 3.904 |  | 1 |  | 3.904 |  | 35.862 |  | < .001 |  | 0.719 |
| Residuals |  | 1.524 |  | 14 |  | 0.109 |  |  |  |  |  |  |
| Interaction |  | 11.005 |  | 1 |  | 11.005 |  | 35.344 |  | < .001 |  | 0.716 |
| Residuals |  | 4.359 |  | 14 |  | 0.311 |  |  |  |  |  |  |
| Condition ✻ Correction |  | 0.023 |  | 1 |  | 0.023 |  | 0.364 |  | 0.556 |  | 0.025 |
| Residuals |  | 0.873 |  | 14 |  | 0.062 |  |  |  |  |  |  |
| Condition ✻ Interaction |  | 0.152 |  | 1 |  | 0.152 |  | 1.667 |  | 0.218 |  | 0.106 |
| Residuals |  | 1.273 |  | 14 |  | 0.091 |  |  |  |  |  |  |
| Correction ✻ Interaction |  | 2.015 |  | 1 |  | 2.015 |  | 30.620 |  | < .001 |  | 0.686 |
| Residuals |  | 0.921 |  | 14 |  | 0.066 |  |  |  |  |  |  |
| Condition ✻ Correction ✻ Interaction |  | 0.276 |  | 1 |  | 0.276 |  | 6.077 |  | 0.027 |  | 0.303 |
| Residuals |  | 0.635 |  | 14 |  | 0.045 |  |  |  |  |  |  |
| *Note.*  Type III Sum of Squares | | | | | | | | | | | | |

| **SUPPLEMENTARY TABLE 11 – Pe amplitude – Mixed ANOVA PD ON vs HC** | | | | | | | | | | | | |
| --- | --- | --- | --- | --- | --- | --- | --- | --- | --- | --- | --- | --- |
| **Within Subjects Effects** | | | | | | | | | | | | |
| **Cases** | | **Sum of Squares** | | **df** | | **Mean Square** | | **F** | | **p** | | **η²_p_** |
| Correction |  | 4.067 |  | 1 |  | 4.067 |  | 61.621 |  | < .001 |  | 0.688 |
| Correction ✻ GROUP |  | 0.037 |  | 1 |  | 0.037 |  | 0.556 |  | 0.462 |  | 0.019 |
| Residuals |  | 1.848 |  | 28 |  | 0.066 |  |  |  |  |  |  |
| Interaction |  | 13.115 |  | 1 |  | 13.115 |  | 53.409 |  | < .001 |  | 0.656 |
| Interaction ✻ GROUP |  | 0.481 |  | 1 |  | 0.481 |  | 1.957 |  | 0.173 |  | 0.065 |
| Residuals |  | 6.876 |  | 28 |  | 0.246 |  |  |  |  |  |  |
| Correction ✻ Interaction |  | 2.181 |  | 1 |  | 2.181 |  | 30.169 |  | < .001 |  | 0.519 |
| Correction ✻ Interaction ✻ GROUP |  | 0.339 |  | 1 |  | 0.339 |  | 4.691 |  | 0.039 |  | 0.143 |
| Residuals |  | 2.024 |  | 28 |  | 0.072 |  |  |  |  |  |  |
| *Note.*  Type III Sum of Squares | | | | | | | | | | | | |

| **Between Subjects Effects** | | | | | | | | | | | | |
| --- | --- | --- | --- | --- | --- | --- | --- | --- | --- | --- | --- | --- |
| **Cases** | | **Sum of Squares** | | **df** | | **Mean Square** | | **F** | | **p** | | **η²_p_** |
| GROUP |  | 0.044 |  | 1 |  | 0.044 |  | 0.233 |  | 0.633 |  | 0.008 |
| Residuals |  | 5.319 |  | 28 |  | 0.190 |  |  |  |  |  |  |
| *Note.*  Type III Sum of Squares | | | | | | | | | | | | |

| **SUPPLEMENTARY TABLE 12 – Pe amplitude – Mixed ANOVA PD OFF vs HC** | | | | | | | | | | | | |
| --- | --- | --- | --- | --- | --- | --- | --- | --- | --- | --- | --- | --- |
| **Within Subjects Effects** | | | | | | | | | | | | |
| **Cases** | | **Sum of Squares** | | **df** | | **Mean Square** | | **F** | | **p** | | **η²_p_** |
| Correction |  | 4.697 |  | 1 |  | 4.697 |  | 62.131 |  | < .001 |  | 0.689 |
| Correction ✻ GROUP |  | 0.002 |  | 1 |  | 0.002 |  | 0.022 |  | 0.883 |  | 7.898×10^-4^ |
| Residuals |  | 2.117 |  | 28 |  | 0.076 |  |  |  |  |  |  |
| Intreraction |  | 16.086 |  | 1 |  | 16.086 |  | 60.514 |  | < .001 |  | 0.684 |
| Intreraction ✻ GROUP |  | 0.092 |  | 1 |  | 0.092 |  | 0.348 |  | 0.560 |  | 0.012 |
| Residuals |  | 7.443 |  | 28 |  | 0.266 |  |  |  |  |  |  |
| Correction ✻ Intreraction |  | 4.007 |  | 1 |  | 4.007 |  | 52.563 |  | < .001 |  | 0.652 |
| Correction ✻ Intreraction ✻ GROUP |  | 0.003 |  | 1 |  | 0.003 |  | 0.043 |  | 0.837 |  | 0.002 |
| Residuals |  | 2.134 |  | 28 |  | 0.076 |  |  |  |  |  |  |
| *Note.*  Type III Sum of Squares | | | | | | | | | | | | |

| **Between Subjects Effects** | | | | | | | | | | | | |
| --- | --- | --- | --- | --- | --- | --- | --- | --- | --- | --- | --- | --- |
| **Cases** | | **Sum of Squares** | | **df** | | **Mean Square** | | **F** | | **p** | | **η²_p_** |
| GROUP |  | 0.013 |  | 1 |  | 0.013 |  | 0.064 |  | 0.802 |  | 0.002 |
| Residuals |  | 5.656 |  | 28 |  | 0.202 |  |  |  |  |  |  |
| *Note.*  Type III Sum of Squares | | | | | | | | | | | | |

| **SUPPLEMENTARY TABLE 13 – Theta induced power – ANOVA PD ON vs OFF** | | | | | | | | | | | | |
| --- | --- | --- | --- | --- | --- | --- | --- | --- | --- | --- | --- | --- |
| **Within Subjects Effects** | | | | | | | | | | | | |
| **Cases** | | **Sum of Squares** | | **df** | | **Mean Square** | | **F** | | **p** | | **η²_p_** |
| Condition |  | 0.093 |  | 1 |  | 0.093 |  | 1.390 |  | 0.258 |  | 0.090 |
| Residuals |  | 0.936 |  | 14 |  | 0.067 |  |  |  |  |  |  |
| Interaction |  | 9.401 |  | 1 |  | 9.401 |  | 40.110 |  | < .001 |  | 0.741 |
| Residuals |  | 3.281 |  | 14 |  | 0.234 |  |  |  |  |  |  |
| Correction |  | 1.296 |  | 1 |  | 1.296 |  | 20.694 |  | < .001 |  | 0.596 |
| Residuals |  | 0.877 |  | 14 |  | 0.063 |  |  |  |  |  |  |
| Condition ✻ Interaction |  | 0.709 |  | 1 |  | 0.709 |  | 7.553 |  | 0.016 |  | 0.350 |
| Residuals |  | 1.315 |  | 14 |  | 0.094 |  |  |  |  |  |  |
| Condition ✻ Correction |  | 0.043 |  | 1 |  | 0.043 |  | 2.081 |  | 0.171 |  | 0.129 |
| Residuals |  | 0.291 |  | 14 |  | 0.021 |  |  |  |  |  |  |
| Interaction ✻ Correction |  | 1.028 |  | 1 |  | 1.028 |  | 19.828 |  | < .001 |  | 0.586 |
| Residuals |  | 0.726 |  | 14 |  | 0.052 |  |  |  |  |  |  |
| Condition ✻ Interaction ✻ Correction |  | 0.009 |  | 1 |  | 0.009 |  | 0.181 |  | 0.677 |  | 0.013 |
| Residuals |  | 0.658 |  | 14 |  | 0.047 |  |  |  |  |  |  |
| *Note.*  Type III Sum of Squares | | | | | | | | | | | | |

| **SUPPLEMENTARY TABLE 14 – Theta induced power – Mixed ANOVA PD ON vs HC** | | | | | | | | | | | | |
| --- | --- | --- | --- | --- | --- | --- | --- | --- | --- | --- | --- | --- |
| **Within Subjects Effects** | | | | | | | | | | | | |
| **Cases** | | **Sum of Squares** | | **df** | | **Mean Square** | | **F** | | **p** | | **η²_p_** |
| Interaction |  | 4.948 |  | 1 |  | 4.948 |  | 45.179 |  | < .001 |  | 0.617 |
| Interaction ✻ GROUP |  | 2.613×10^-7^ |  | 1 |  | 2.613×10^-7^ |  | 2.385×10^-6^ |  | 0.999 |  | 8.519×10^-8^ |
| Residuals |  | 3.067 |  | 28 |  | 0.110 |  |  |  |  |  |  |
| Correction |  | 2.358 |  | 1 |  | 2.358 |  | 58.946 |  | < .001 |  | 0.678 |
| Correction ✻ GROUP |  | 0.036 |  | 1 |  | 0.036 |  | 0.895 |  | 0.352 |  | 0.031 |
| Residuals |  | 1.120 |  | 28 |  | 0.040 |  |  |  |  |  |  |
| Interaction ✻ Correction |  | 0.956 |  | 1 |  | 0.956 |  | 24.793 |  | < .001 |  | 0.470 |
| Interaction ✻ Correction ✻ GROUP |  | 0.016 |  | 1 |  | 0.016 |  | 0.425 |  | 0.520 |  | 0.015 |
| Residuals |  | 1.080 |  | 28 |  | 0.039 |  |  |  |  |  |  |
| *Note.*  Type III Sum of Squares | | | | | | | | | | | | |

| **Between Subjects Effects** | | | | | | | | | | | | |
| --- | --- | --- | --- | --- | --- | --- | --- | --- | --- | --- | --- | --- |
| **Cases** | | **Sum of Squares** | | **df** | | **Mean Square** | | **F** | | **p** | | **η²_p_** |
| GROUP |  | 0.330 |  | 1 |  | 0.330 |  | 1.619 |  | 0.214 |  | 0.055 |
| Residuals |  | 5.709 |  | 28 |  | 0.204 |  |  |  |  |  |  |
| *Note.*  Type III Sum of Squares | | | | | | | | | | | | |

| **SUPPLEMENTARY TABLE 15 – Theta induced power – Mixed ANOVA PD OFF vs HC** | | | | | | | | | | | | |
| --- | --- | --- | --- | --- | --- | --- | --- | --- | --- | --- | --- | --- |
| **Within Subjects Effects** | | | | | | | | | | | | |
| **Cases** | | **Sum of Squares** | | **df** | | **Mean Square** | | **F** | | **p** | | **η²_p_** |
| Interaction |  | 9.404 |  | 1 |  | 9.404 |  | 63.682 |  | < .001 |  | 0.695 |
| Interaction ✻ GROUP |  | 0.708 |  | 1 |  | 0.708 |  | 4.797 |  | 0.037 |  | 0.146 |
| Residuals |  | 4.135 |  | 28 |  | 0.148 |  |  |  |  |  |  |
| Correction |  | 1.762 |  | 1 |  | 1.762 |  | 57.288 |  | < .001 |  | 0.672 |
| Correction ✻ GROUP |  | 0.158 |  | 1 |  | 0.158 |  | 5.128 |  | 0.031 |  | 0.155 |
| Residuals |  | 0.861 |  | 28 |  | 0.031 |  |  |  |  |  |  |
| Interaction ✻ Correction |  | 0.784 |  | 1 |  | 0.784 |  | 13.566 |  | < .001 |  | 0.326 |
| Interaction ✻ Correction ✻ GROUP |  | 0.001 |  | 1 |  | 0.001 |  | 0.022 |  | 0.883 |  | 7.878×10^-4^ |
| Residuals |  | 1.619 |  | 28 |  | 0.058 |  |  |  |  |  |  |
| *Note.*  Type III Sum of Squares | | | | | | | | | | | | |

| **Between Subjects Effects** | | | | | | | | | | | | |
| --- | --- | --- | --- | --- | --- | --- | --- | --- | --- | --- | --- | --- |
| **Cases** | | **Sum of Squares** | | **df** | | **Mean Square** | | **F** | | **p** | | **η²_p_** |
| GROUP |  | 0.073 |  | 1 |  | 0.073 |  | 0.358 |  | 0.555 |  | 0.013 |
| Residuals |  | 5.700 |  | 28 |  | 0.204 |  |  |  |  |  |  |
| *Note.*  Type III Sum of Squares | | | | | | | | | | | | |

| **Supplemetary Table 16 - Linear Mixed Models results** | | | | | | |
| --- | --- | --- | --- | --- | --- | --- |
| Type III Analysis of Variance Table with Satterthwaite's method | | | | | | |
|  | **Sum Sq** | **Mean Sq** | **NumDF** | **DenDF** | **F Value** | **Pr(>F)** |
| Group | 2217719 | 1108860 | 2 | 53.9 | 32.6642 | 5.126e-10 * |
| Condition | 112027015 | 112027015 | 1 | 9491.1 | 3300.0326 | < 2.2e-16 *** |
| Theta | 114 | 114 | 1 | 9492.6 | 0.0034 | 0.953709 |
| Group:Condition | 10614054 | 5307027 | 2 | 9491.1 | 156.3316 | < 2.2e-16 *** |
| Group:Theta | 196627 | 98313 | 2 | 9492.1 | 2.8961 | 0.055289 |
| Condition:Theta | 282501 | 282501 | 1 | 9492.8 | 8.3218 | 0.003926 |
| Group:Condition:Theta | 24992 | 12496 | 2 | 9492.5 | 0.3681 | 0.692061 |

**Supplementary Table 17 | Results of Neuropsychological Tests of PD patients.**

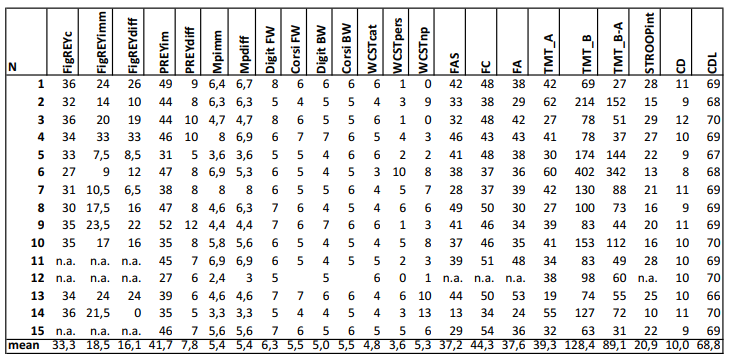

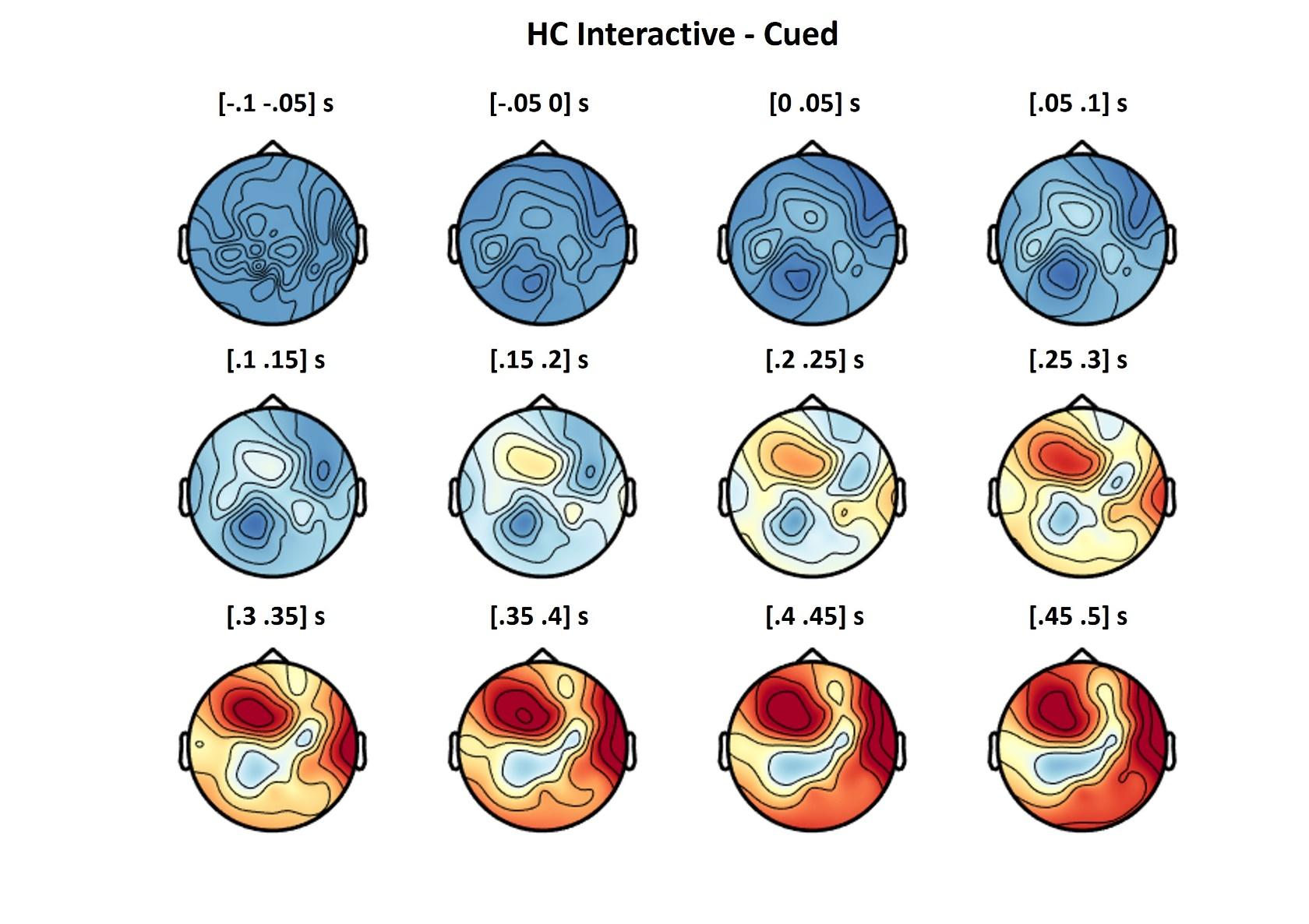

**Figure S1 |** Topographies of the activity in the Theta band over time, as shown in Fig. 6, for the HC group.

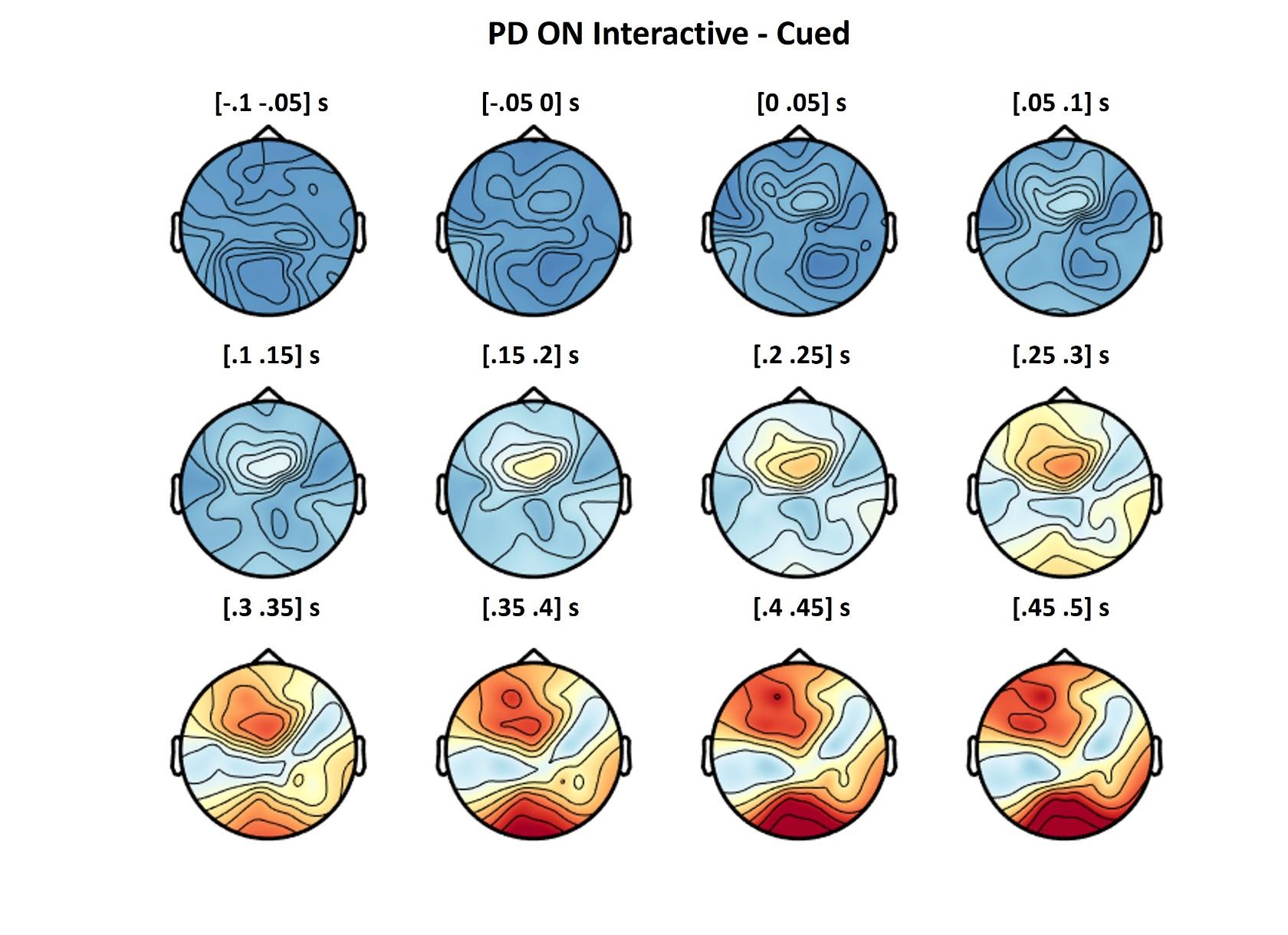
**Figure S2 |** Topographies of the activity in the Theta band over time, as shown in Fig. 6, for the PD ON group.

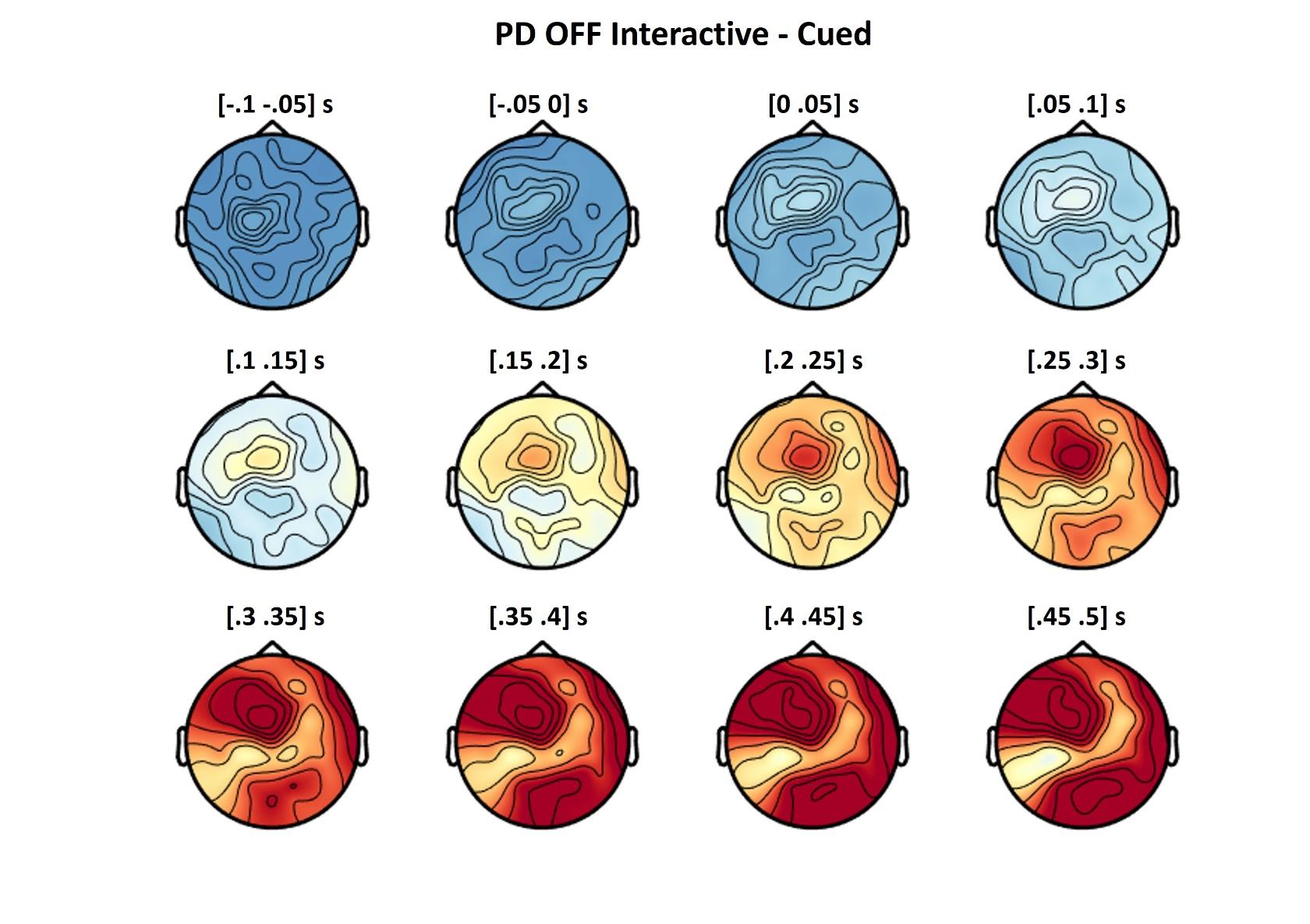

**Figure S3 |** Topographies of the activity in the Theta band over time, as shown in Fig. 6, for the PD OFF group.

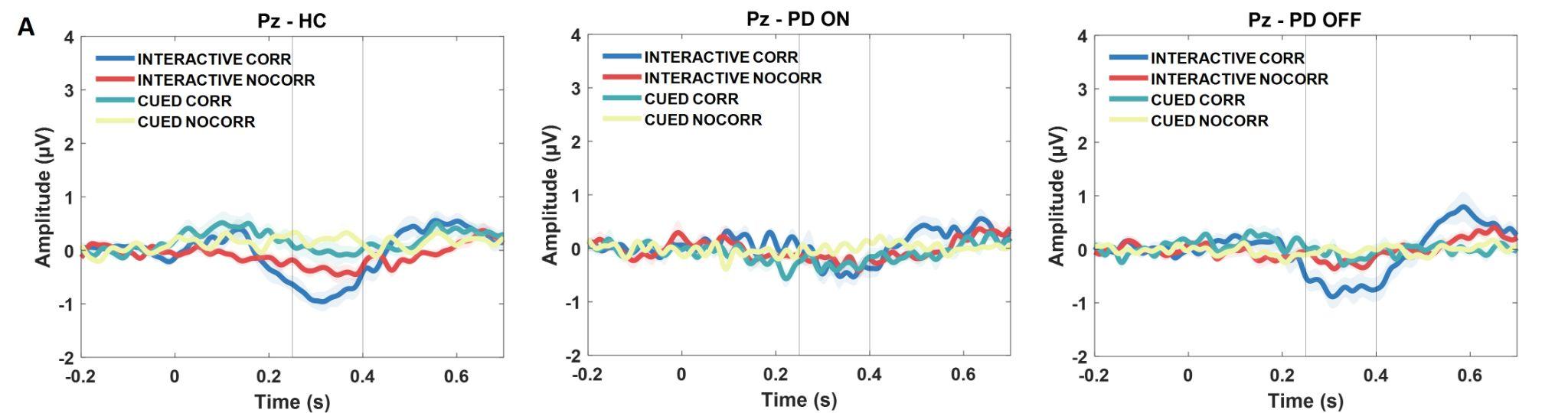

**Figure S4 |** Grand averages of the Pe component over Pz in all experimental conditions, divided by group (HC, PD ON/OFF). Data are time-locked to the correction of the Virtual Partner (or the equivalent frame when no correction occurred).

**Supplementary references**

Albert, N. B., Peiris, Y., Cohen, G., Miall, R. C., & Praamstra, P. (2010). Interference effects from observed movement in Parkinson's disease. Journal of Motor Behaviour, 42(2), 145– 149.

Bek, J., Gowen, E., Vogt, S., Crawford, T., & Poliakoff, E. (2018). Action observation produces motor resonance in Parkinson's disease. Journal of neuropsychology, 12(2), 298-311.

Capasso, R., & Miceli, G. (2001). Esame Neuropsicologico dell’Afasia. Springer-Verlag, Milan, Italy.

Carlesimo, G. A., Buccione, I., Fadda, L., Graceffa, A., Mauri, M., Lorusso, S., Caltagirone, C. (2002). Standardizzazione di due test di memoria per uso clinico: breve racconto e figura di Rey. *Nuova Rivista di Neurologia, 12,* 1–13.

Carlesimo, G. A., Caltagirone, C., & Gainotti G. (1996). The Mental Deterioration Battery: normative data, diagnostic reliability and qualitative analyses of cognitive impairment. The Group for the Standardization of the Mental Deterioration Battery. *European Neurology, 36*, 378–384. https://doi.org/[10.1159/000117297](http://dx.doi.org/10.1159/000117297)

Castiello, U., & Bennett, K. M. (1997). The bilateral reach-to-grasp movement of Parkinson's disease subjects. Brain: a journal of neurology, 120(4), 593-604.

Costa, A., Bagoj, E., Monaco, M., Zabberoni, S., De Rosa, S., Papantonio, A. M., Carlesimo, G.A. (2014). Standardization and normative data obtained in the Italian population for a new verbal fluency instrument, the phonemic/semantic alternate fluency test. *Neurological Sciences, 35*, 365, https://doi.org/[10.1007/s10072-013-1520-8](http://dx.doi.org/10.1007/s10072-013-1520-8)

Costa, A., Monaco, M., Zabberoni, S., Peppe, A., Perri, R., Fadda, L., . . . Carlesimo, G. A. (2014). Free and cued recall memory in Parkinson’s disease associated with amnestic mild cognitive impairment. *PLoS One, 9,* e86233. <https://doi.org/10.1371/journal.pone.0086233>

Fahn, S., & Elton, R. L. (1987). “*UPDRS development committee. Unified Parkinson’s disease rating scale,*” in Recent Development in Parkinson’s Disease, eds S. Fahn, P. D. Marsden, D. B. Calne, and A. Liebarman (Florman Park, NJ: Mac Millan Health Care Information), 153–163.

Giovagnoli, A. R., Del Pesce, M., Mascheroni, S., Simoncelli, M., Laiacona, M., & Capitani, E. (1996). Trail making test: normative values from 287 normal adult controls. *Italian Journal of Neurological Sciences, 17*, 305–309. <https://doi.org/10.1007/BF01997792>

Hoehn, M. M., & Yahr, M. D. (1967). Parkinsonism: Onset, Progression and Mortality. *Neurology, 17*, 427-42. <https://doi.org/10.1212/wnl.17.5.427>

Lukos, J. R., Snider, J., Hernandez, M. E., Tunik, E., Hillyard, S., & Poizner, H. (2013). Parkinson’s disease patients show impaired corrective grasp control and eye–hand coupling when reaching to grasp virtual objects. Neuroscience, 254, 205-221.

Measso, G., Cavarzeran, F., Zappala, G., Lebowitz, B. D., Crook, T. H., Pirozzolo, F .J., . . . Grigoletto, F. (1991). The Mini mental state Examination: normative study of a random sample of the Italian population. *Developmental Neuropsychology, 9*, 77–85. <https://doi.org/10.1080/87565649109540545>

Monaco, M., Costa, A., Caltagirone, C., & Carlesimo, G. A. (2013). Forward and backward span for verbal and visuo-spatial data: standardization and normative data from an Italian adult population. *Neurological Sciences, 34*, 749–754. [https://doi.org/](https://doi.org/10.1002/mds.21507)[10.1007/s10072-012-1130-x](http://dx.doi.org/10.1007/s10072-012-1130-x)

Nocentini, U., Di Vincenzo, S., Panella, M., Pasqualetti, P., & Caltagirone, C. (2002). La valutazione delle funzioni esecutive nella pratica neuropsicologica; dal modified Card sorting test al modified card sorting test-Roma version. Dati di standardizzazione. *Nuova Rivista di Neurologia, 12*, 13–24.

Poliakoff, E., Galpin, A., Dick, J., Moore, P., & Tipper, S. P. (2007). The effect of viewing graspable objects and actions in Parkinson's disease. NeuroReport, 18, 483– 487.
